# Targeting melanosome pH is an effective method for the treatment of oculocutaneous albinism

**DOI:** 10.64898/2026.05.25.727673

**Authors:** Samuel J Grondin, Daniela St. Pierre, David J. Green, Sidra Amir, Maftuna Yusupova, Joseph Bonica, Zuhal Eraslan, Tyler Wills, Callum Hunt, Dalee Zhou, Aman George, Jaewon You, Aditya Anandakumar, Steven S Gross, Ryan Schreiner, Qiuying Chen, Mervyn G. Thomas, Stacie K. Loftus, David R. Adams, Kazumasa Wakamatsu, Shosuke Ito, Panagiotis I. Sergouniotis, Melissa Harris, Brian P. Brooks, Jonathan H. Zippin

## Abstract

Oculocutaneous albinism (OCA) is a genetic condition associated with impaired visual acuity and increased skin cancer risk. When OCA is due to defects in melanosome ion transport, abnormally acidic conditions in the melanosome lumen inhibit tyrosinase, the critical pigment synthetic enzyme. Hence, a therapeutic approach that optimizes melanosome pH to increase pigment production presents a potential treatment for OCA and a method for decreasing skin cancer risk. Here, we report that reduction in sAC (*ADCY10*) activity via naturally occurring human variants in *ADCY10* restores OCA pigmentation, and sAC inhibition increases melanin synthesis in both human and mouse OCA models. These findings demonstrate that targeting melanosome pH is an effective, previously untapped therapeutic strategy for OCA and elevated skin cancer risk.

## Introduction

Oculocutaneous albinism (OCA) is an autosomal recessive disease characterized by reduced pigmentation in the skin, hair and eyes(*1*) that affects >300,000 people globally(*2*). Decreased skin pigmentation leads to a markedly increased risk of all forms of skin cancer, and without UV protection, many affected individuals develop metastatic disease at a young age(*2, 3*). OCA is also associated with nystagmus, foveal hypoplasia, and other ocular manifestations that cause visual impairment(*1*). As an orphan disease, there is a dearth of therapeutic options to correct the underlying pathology, and management remains largely symptomatic.

In mammals there are two types of cells that synthesize melanin: pigmented epithelial cells in the eye (e.g., retinal pigmented epithelial cells (RPE)) and melanocytes in the skin, eye (iris, ciliary body and choroid), and various extracutaneous sites(*4, 5*). There are at least eight types of non-syndromic OCA defined by mutations leading to reduced pigment production in these cell types(*6*). Three of them result from reduced activity of core melanin biosynthetic enzymes (*TYR*, *TYRP1*, *DCT*)(*6*). The remaining forms arise primarily from defects in membrane ion channels and transporters (including *OCA2*, *SLC45A2* and *SLC24A5*), which are thought to acidify the lumen of the melanosome, the specialized melanin producing organelle(*6–9*). Increased acidity lowers the activity of tyrosinase (TYR), the rate-limiting enzyme in melanin synthesis(*10, 11*). Conversely, alkalinizing melanosome pH to a near-neutral pH enhances TYR activity and, in principle, could correct pigment deficits across most forms of OCA(*10, 11*).

Many individuals in the general population carry commonly occurring genetic variants that alter the function of melanosome ion channels linked to OCA(*12*). Such variants shift melanosome pH to a more acidic setpoint which is thought to contribute to observed fair skin pigmentation and increased skin cancer risk. These population level effects highlight the broad clinical relevance of impaired melanosome pH. Previous therapeutic efforts directed at improving pigmentation have largely focused on stimulating TYR directly rather than correcting melanosome pH. Relevant approaches have included supplying high levels of melanin metabolites to bypass TYR activity (e.g., L-DOPA)(*13*) or increasing tyrosine availability (e.g., nitisinone) to force additional TYR activity(*14*). These strategies have shown limited efficacy, possibly because the pH of the melanosome is the rate-limiting factor and is independent of the amount of TYR present. Gene therapeutic approaches, such as adeno-associated virus-mediated transfer of a functional *TYR* cDNA or CRISPR/Cas9-mediated correction of *TYR* deleterious variants, have shown some promise in animal models, particularly for localized therapeutic areas (e.g., eye), but are unlikely to be effective for widespread, multi-organ disease(*15*). To date, there are no therapeutic approaches designed to rescue melanin synthesis by addressing melanosome pH.

Our previous work demonstrated that inhibition of sAC alkalinizes melanosome pH in OCA melanocytes(*16*), suggesting sAC inhibitors could restore melanin synthesis in OCA. Here, we demonstrate that reducing sAC activity (genetically or pharmacologically) restores pigmentation in human and mouse models of OCA.

## Results

### Genetic variants predicted to reduce *sAC* (ADCY10) function are associated with darker skin and hair pigmentation in the presence or absence of OCA2 or TYR deleterious variants

We analysed UK Biobank participants carrying predicted deleterious variants in *ADCY10* (the gene encoding sAC), *OCA2* (the gene responsible for OCA type 2), and *TYR* (the gene responsible for OCA type 1). As expected, *OCA2* variants associated with reduced function were strongly linked to lighter skin and hair pigmentation (Fig. 1A, tables S1,3). In contrast, variants predicted to reduce *ADCY10* function were associated with darker pigmentation (Fig. 1A, tables S1,3). Notably, individuals carrying variants in both genes exhibited pigmentation indistinguishable from those carrying *ADCY10* variants alone (Fig. 1A, tables S1,3). Similarly, *TYR* variants were associated with reduced skin and hair pigmentation (Fig. 1B, tables S2,4) while individuals carrying variants in both *TYR* and *ADCY10* were found to have substantially restored pigmentation of the hair and skin (that exceeded that of the reference alleles) (Fig. 1B, tables S2,4). These observations suggested that reduced *ADCY10* activity could counteract the hypopigmenting effects of partial *OCA2* or *TYR* deficiency, providing population-level evidence consistent with the hypothesis that sAC inhibition could restore melanin synthesis in OCA melanocytes and RPE cells.

**Fig. 1.**
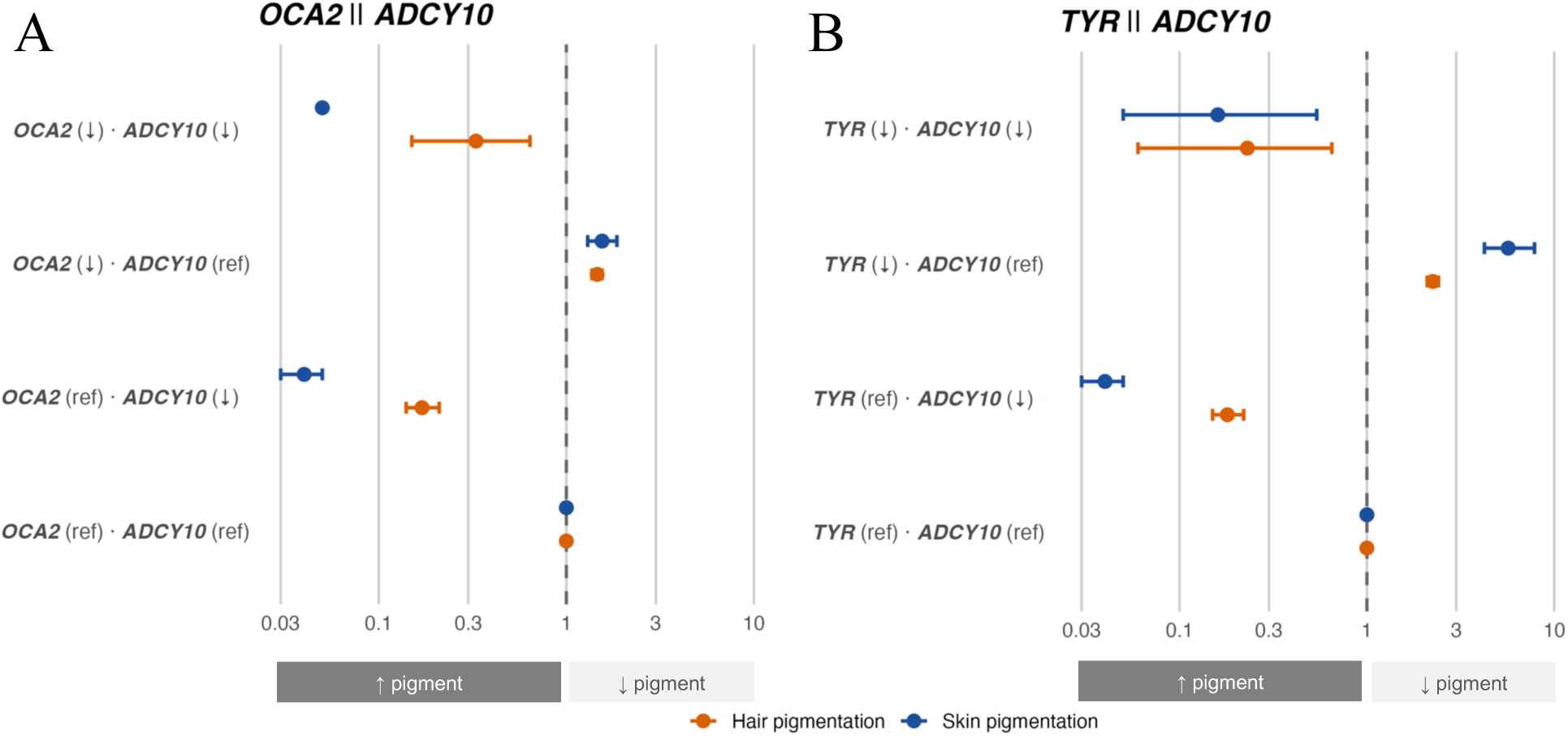
Deleterious variants in *ADCY10* associate with darker skin and hair pigmentation, including in the context of *OCA2* or *TYR* partial loss of function. Forest plots showing odds ratios from binomial logistic regression models testing the association between genotype groups and pigmentation phenotypes for *OCA2*–*ADCY10* (A) and *TYR*–*ADCY10* (B). Genotype groups were defined as described in the Methods. Odds ratios represent the odds of light relative to dark pigmentation, with genotype group A used as the reference category. Points indicate odds ratios and horizontal bars indicate 95% confidence intervals; the dashed vertical line denotes an odds ratio of 1. Analyses were performed separately for hair (orange) and skin (dark blue) pigmentation. Group sizes for each analysis and relevant numerical data are provided in tables S1-4. ↓ denotes reduced gene function due to the presence of ≥1 functional variant; “ref” denotes absence of a functional variant.

### Alkalinization of melanosome pH increases melanin synthesis in pigmented cells from individuals with OCA

As mentioned, abnormally acidic melanosome pH is the presumed pathophysiology of many types of OCA(*6, 7, 9*). Thus, we hypothesized that alkalinization of melanosome pH might improve melanin synthesis in individuals with OCA subtypes. Traditional methods for alkalinizing melanosome pH rely on inhibitors of proton pumps (e.g., bafilomycin) which are effective but not suitable for therapeutic use due to toxicity(*11, 17*). Recently improved sAC inhibitors (sACi) were developed which have nanomolar IC_50_, no observable toxicity(*18–21*), and are effective at alkalinizing melanosome pH and increasing melanin synthesis in normal melanocytes(*16*). sACi are safe and effective at blocking sAC activity for extended periods of time in mouse models(*19*) and have passed multiple *in vitro* and *in vivo* tests for drug safety(*20, 21*).

Our group and others have reported that melanosome pH is more acidic in pigmented cells lacking OCA2 activity(*16, 22*). We previously demonstrated that sACi treatment alkalinized melanosome pH in human melanocytes(*23*) and we now show that sACi treatment alkalinized melanosome pH in human RPE cells (fig. S1). Thus, sAC inhibition is an effective method for alkalinizing melanosome pH in all types of mammalian pigmented cells. Moreover, we recently reported that sACi treatment rescued abnormally acidic melanosome pH back to wild type levels in a variety of OCA model melanocytes (e.g., OCA type 2, OCA type 4, OA1)(*16*) and we now show that sACi treatment increased TYR activity in live *Oca2* null cells within hours (fig. S2A), consistent with sAC inhibition increasing melanin synthesis by augmenting the activity of existing TYR.

Human primary melanocytes derived from individuals with OCA type 2 resulting from multiple distinct *OCA2* variants (p.V443I/p.V443I or D2.7kb/D2.7kb) were incubated with sACi for 72 hours. We observed a significant increase in dark granules within OCA type 2 melanocytes as compared to vehicle treated cells (Fig. 2A-B and fig. S2B-C). Using our recently reported flow cytometry method, we confirmed that sAC inhibition of human and mouse OCA type 2 melanocytes led to increased scatter of 355 nM light consistent with increased melanin synthesis (Fig. 2C, fig. S2D) (*24*). Ultrastructural analysis of human OCA type 2 melanocytes treated with vehicle revealed mostly poorly-melanized melanosomes and very few highly-melanized melanosomes, likely due to low levels of overall melanin synthesis in these cells (Fig. 2D,F). In contrast, treatment with sACi led to a significant increase in highly-melanized melanosomes and a concomitant decrease in poorly-melanized melanosomes (Fig. 2E-F). To confirm that the increase in highly-melanized melanosomes was not due to an overall increase in the production of melanosomes, we performed immunocytochemistry using melanosome markers. In human and mouse OCA type 2 melanocytes, treatment with sACi did not affect the total number of melanosomes positive for HMB45 (early-stage melanosome marker) or TYRP1 (late-stage melanosome marker) suggesting that sAC inhibition does not overtly affect melanosome genesis over 72 hours (fig. S3A-D).

**Fig. 2.**
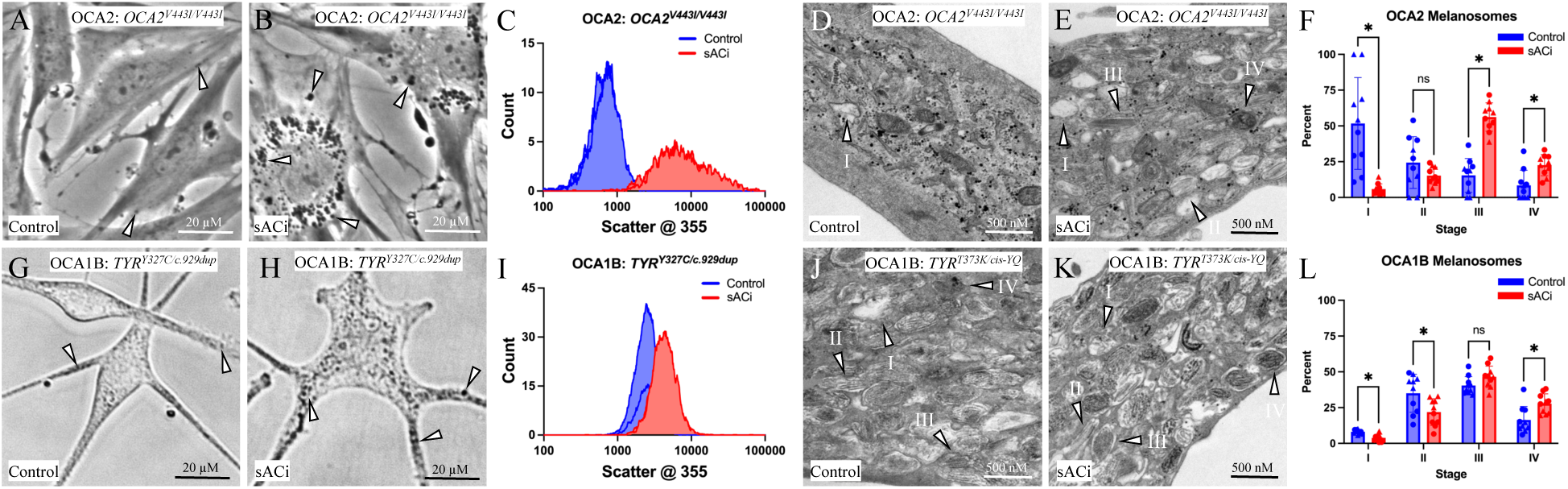
Inhibition of sAC increases melanin synthesis in melanocytes from patients with OCA2 and OCA1B. A-B) Human melanocytes derived from a patient with OCA2 (*V443I/V443I*) were treated with vehicle (DMSO) or sACi (TDI-11155, 30µM) for 24 hours. sACi treatment increased the number of dark colored organelles (arrows) by phase contrast microscopy. Representative images from an experiment performed N=3. C) Flow cytometry of the cells shown in A-B revealed an increase in scatter at 355 nm consistent with increased melanin synthesis. N=3. D-E) Electron microscopy of the same conditions described in A-B revealed an increase in the number of late-stage melanosomes. Representative images from an experiment performed N=3. F) Percent distribution of melanosomes at different stages quantitated from at least 20 electron micrographs consisting of at least five distinct melanocytes derived from two distinct OCA2 patient samples (circles vs triangles). G-H) Human melanocytes derived from a patient with OCA1B (*TYR^Y327C/c.929dup^*) were treated with vehicle (DMSO) or sACi (TDI-11155, 30µM) for 24 hours. sACi treatment increased the number of dark colored organelles (arrows) by phase contrast microscopy. I) Flow cytometry of the cells shown in G-H revealed an increase in scatter at 355 nm consistent with increased melanin synthesis. J-K) Electron microscopy of the same conditions described in G-H revealed an increase the number of late-stage melanosomes. Representative images from an experiment performed N=3. L) Percent distribution of melanosomes at different stages quantitated from at least 20 electron micrographs consisting of at least five distinct melanocytes derived from two distinct OCA1B patient samples (circles vs triangles). F, L) Multiple Welch’s t-test with Holm-Šídák correction. *, P<0.05.

As OCA type 2 also affects pigmentation in the eye, we sought to examine the effects of sACi on RPE pigmentation. We generated human RPE cells with normal or *OCA2* loss of function by differentiating CRISPR gene edited induced pluripotent stem cells (iPSC) cells into RPE cells(*25, 26*). Treatment with sACi increased melanin content in wild type human RPE cells (fig. S2E), confirming that alkalinization of melanosome pH in wild type human ARPE-19 cells (fig. S1) can increase melanin synthesis in these cells. Similar to OCA type 2 melanocytes, treatment with sACi increased melanin synthesis in OCA type 2 RPE cells as measured by flow cytometry (fig. S2F). Thus, sAC inhibition increased melanin synthesis in both melanocytes and RPE cells with OCA2 loss of function.

OCA type 4 has a similar clinical presentation to OCA type 2 but arises from loss of function of a different melanosome ion channel, SLC45A2, which, like OCA2 loss of function, leads to a more acidic melanosome pH(*9*). As observed in OCA type 2 melanocytes, sAC inhibition of SLC45A2 knockout mouse melanocytes also increased melanin synthesis (fig. S2G-J and table S5). Since alkalinization of melanosome pH increases TYR activity under multiple conditions, we reasoned sAC inhibition might increase melanin synthesis in any type of OCA where the TYR enzyme retains some activity. Thus, we next asked if sAC inhibition elevated melanin synthesis in patient-derived OCA type 1B human melanocytes which contain a hypomorphic tyrosinase enzyme. Treatment of multiple OCA type 1B human melanocyte lines with distinct causative genotypes (p.T327C/p.R311X and p.T373K/p.[S192Y;R402Q]) with sACi increased melanin synthesis as measured visually and by flow cytometry (Fig. 2G-I). In addition, ultrastructural analysis of OCA type 1B melanocytes treated with sACi showed significantly more highly-melanized melanosomes (Fig. 2J-L). As in OCA type 2 melanocytes, increased melanin synthesis following sACi treatment was not associated with any changes in total melanosome number or markers of melanogenesis (fig. S3E-F). Taken together, these data suggest that sAC inhibition increases pigmentation in multiple patient-derived OCA models without increasing melanosome genesis.

### Genetic deletion of sAC increases ocular and cutaneous pigmentation in an OCA2 mouse model

Since sACi treatment increased melanin synthesis in isolated OCA human and mouse melanocytes and RPE cells, we reasoned that sAC inhibition might rescue tissue pigmentation in an albino mouse model. Therefore, we assessed ocular and cutaneous pigmentation following genetic deletion of sAC in *Tyr* expressing cells in an OCA type 2 mouse model. We previously developed a mouse model which utilizes the *Tyr::Cre^ERT2^* allele to allow for the inducible knockout of sAC (*Adcy10^fl/fl^*) exclusively in cells expressing the tyrosinase gene (melanocytes and RPE)(*23, 27*). We introduced an *Oca2* null allele (*28*) to homogeneity and backcrossed this mouse onto a C3H/HeJ agouti mouse strain for five generations to enable the assessment of both eumelanin and pheomelanin synthesis, producing a mouse with the complete genotype C3H/HeJ; *Tyr::Cre^ERT2^*; *Adcy10^fl/fl^*; *Oca2^p.R262X/p.R262X^*(*sAC^fl/fl^; OCA2^-/-^*) (Fig. 3A and fig S4). In all instances *Tyr::Cre^ERT2^* + and – mice were mated to each other, and the entire litter was topically treated across the entire skin and eyes with 4-hydroxytamoxifen (TAM) on P2-4 (Fig. 3B). This approach allowed us to compare *sAC^-/-^* and *sAC^fl/fl^*littermates. OCA2 null mice with genetic deletion of sAC exhibited an obvious increase in ocular pigmentation (Fig. 3). Optical imaging of the eye revealed an increase in iris and fundus pigmentation at P30 in *sAC^-/-^* compared to *sAC^fl/fl^*littermates (Fig. 3C-F). Microscopic examination demonstrated an increase in melanin levels by Fontana-Masson staining in melanocytes of the iris, ciliary body, and choroid and pigmented epithelial cells of the ciliary body and choroid (e.g., RPE) of the eyes in *sAC^-/-^*animals compared to *sAC^fl/fl^* littermates with no clear increase in pigment gene expression (Fig. 3G-L and fig. S5A-E). Ultrastructural examination revealed that *sAC^-/-^* pigmented cells of the eye had significantly more late-stage melanosomes (Fig. 3M-R and fig. S5F).

**Fig. 3.**
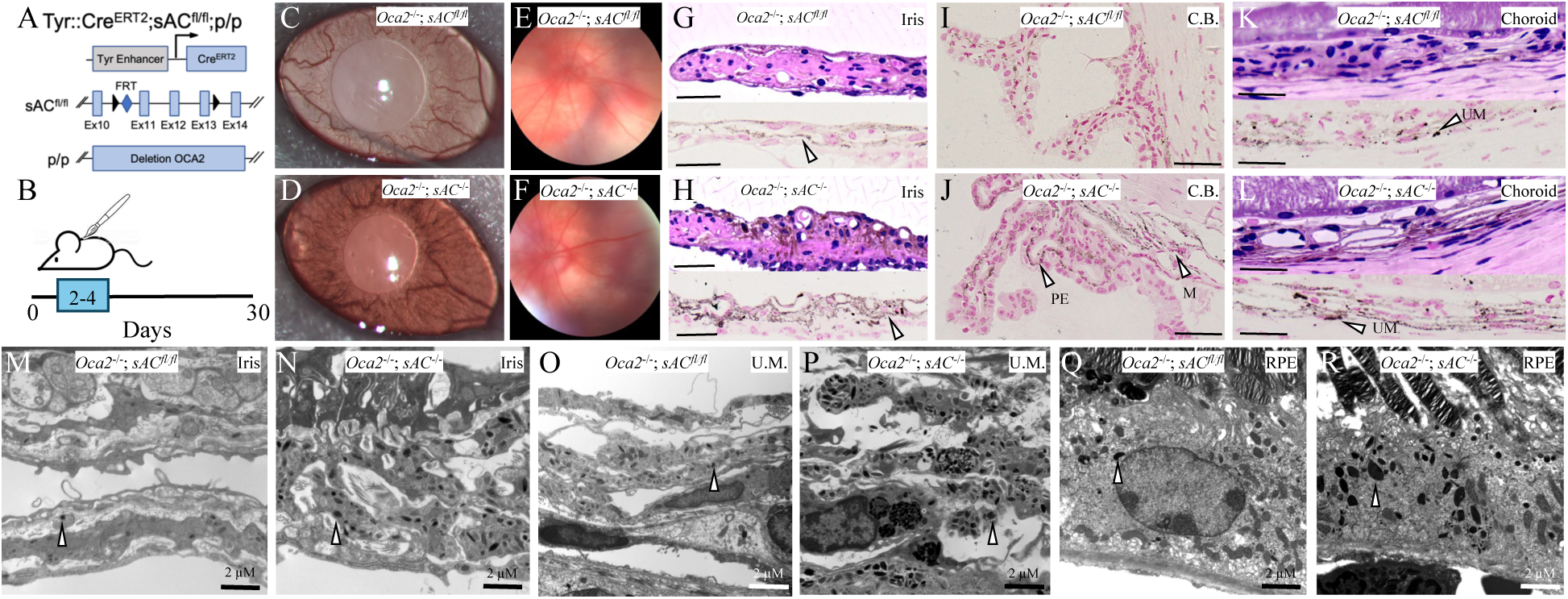
Genetic loss of function of sAC rescues ocular pigmentation in a OCA2 null mouse model. A) Scheme for tamoxifen-induced genetic deletion of sAC in TYR expressing cells in a p/p (*OCA2^-/-^*) mouse model. B) Diagram showing the painting of *Tyr::Cre^ERT2^+ or -; sAC^fl/fl^; p/p* littermates with 4-hydroxytamoxifen on P2-4 with ocular pigmentation observed on P30. C-D) Iris slit lamp images from *sAC^fl/fl^* (C) vs *sAC^-/-^* (D) mice showing darker iris tissue in *sAC^-/-^* mice. E-F) Fundoscopic images from *sAC^fl/fl^* (E) vs *sAC^-/-^* (F) mice showing a darker image in *sAC^-/-^*mice. G-H) Histology of iris from *sAC^fl/fl^* (G) vs *sAC^-/-^* (H) mice stained with H/E (upper panel) or Fontana-Masson (FM, lower panel) showing more FM staining in the iris tissue in *sAC^-/-^* mice. Arrows denote FM+ areas. I-J) FM staining of ciliary body from *sAC^fl/fl^* (I) vs *sAC^-/-^* (J) mice showing increased FM staining in both pigmented epithelial cells (PE) and melanocytes (M) in *sAC^-/-^* mice. Arrows denote FM+ areas. K-L) Histology of choroid from *sAC^fl/fl^* (K) vs *sAC^-/-^* (L) mice stained with H/E (upper panel) or FM (lower panel) showing more FM staining in the choroid in *sAC^-/-^* mice. Arrows denote FM+ areas. UM, uveal melanocytes. M-R) Electron micrographs of whole eyes from *sAC^fl/fl^* (M, O, Q) vs *sAC^-/-^* (N, P, R) mice depicting iris melanocytes (M-N), uveal melanocytes (U.M., O-P), and RPE cells (Q-R) showing an increase in late-stage melanosomes in *sAC^-/-^* mice. Arrows denote stage IV melanosomes. All images are representative of N≥10 mice per cohort, mice combined over at least three litters.

Further, we compared hair color at P30 and found that *sAC^-/-^*mice had visibly darker dorsal and ventral hair compared to *sAC^fl/fl^* animals (Fig. 4A-B and fig. S6A-B). Quantification of melanin by HPLC revealed a significant increase in total hair shaft melanin and eumelanin content (Fig. 4C and fig. S6C-D). Histologic examination of individual hair follicles revealed a mix of both eumelanin- and pheomelanin-synthesizing hair bulbs, as expected in an agouti mouse strain, with *sAC^-/-^* mice exhibiting an overall increase in the percent of eumelanin-positive and a decrease in the percent of pheomelanin-positive follicles (Fig. 4D-E and fig S6E). The melanocyte marker DCT identifies hair follicles with the capacity to synthesize eumelanin. We found that many hair bulbs in *sAC^fl/fl^* animals express DCT but have low or no eumelanin, likely due to OCA2 deficiency, whereas hair bulbs in *sAC^-/-^*animals showed a significant increase in the percent of medium to high eumelanin-producing, DCT+ hair bulbs (Fig. 4F and fig. S6F). SOX10 stains both eumelanin- and pheomelanin-producing melanocytes and was only modestly increased in *sAC^-/-^*mice (fig. S6G-I). These data suggest that melanocytes in *sAC^fl/fl^* animals are primed to produce eumelanin but do not do so in abnormally low melanosome pH caused by OCA type 2. Upon restoration of the melanosome pH by genetic sAC knockout, these hair bulbs resume production of eumelanin. Overall, these observations are consistent with our previously published studies of sAC loss-of-function in wild type agouti mice(*23*) and mirror our observed changes in human melanocyte and mouse hair melanin levels (Figs. 2, 4A-C and fig S6A-D).

**Fig. 4.**
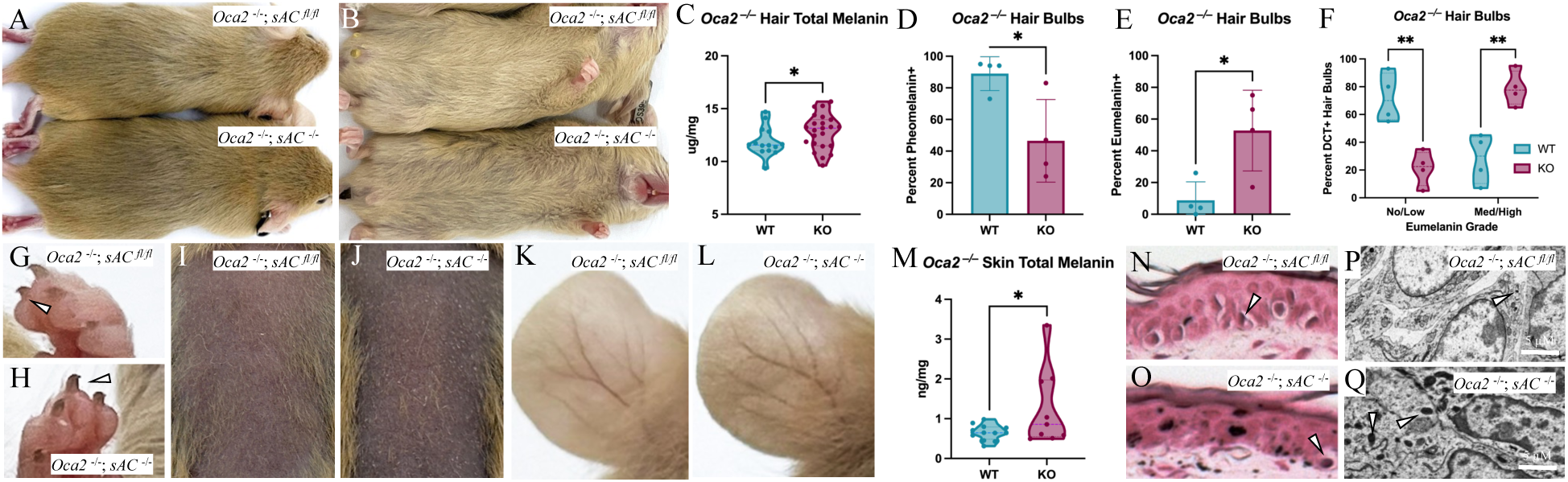
Genetic loss of function of sAC rescues cutaneous pigmentation in a OCA2 null mouse model. A) Dorsal hair from *sAC^fl/fl^* (upper) vs *sAC^-/-^* (lower) mice showing darker hair in *sAC^-/-^* mice. B) Ventral hair from *sAC^fl/fl^* (upper) vs *sAC^-/-^* (lower) mice showing darker hair in *sAC^-/-^* mice. C) HPLC analysis of melanin content of hair from *sAC^fl/fl^* (WT) vs *sAC^-/-^* (KO) mice. Each point is a mouse. N≥14 per cohort. Welch’s t-test, *,P<0.05. D-E) Percent of observed pheomelanin (D) and eumelanin (E) positive hair bulbs in dorsal skin sections from *sAC^fl/fl^* (WT) vs *sAC^-/-^* (KO) mice showing a significant decrease in pheomelanin and increase in eumelanin positive hair bulbs. Welch’s t-test, *P<0.05. Each point is average per mouse. F) Comparison of the change in the number of no/low eumelanin and med/high eumelanin DCT+ hair bulbs in *sAC^fl/fl^* (WT) vs *sAC^-/-^* (KO) mice showing that KO mice possess fewer no/low eumelanin DCT+ hair bulbs and more med/high eumelanin DCT+ hair bulbs than WT mice. Each point is average per mouse. Two-way ANOVA, **P<0.01. G-H) Nails from *sAC^fl/fl^* (G) vs *sAC^-/-^* (H) mice showing darker nails in *sAC^-/-^* mice. Arrows denote nail pigmentation. I-J) Waxed skin from *K14::Kitl^+^ sAC^fl/fl^* (I) vs *K14::Kitl^+^ sAC^-/-^* (J) mice showing darker skin in *sAC^-/-^* mice. K-L) Ears from *K14::Kitl^+^ sAC^fl/fl^* (K) vs *K14::Kitl^+^ sAC^-/-^* (L) mice showing darker skin and hair in *sAC^-/-^*mice. M) HPLC analysis of melanin content in skin from *sAC^fl/fl^* (WT) vs *sAC^-/-^* (KO) mice. N≥9 per cohort. N-O) Fontana-Masson (FM) staining of ear skin from *K14::Kitl^+^ sAC^fl/fl^* (N) vs *K14::Kitl^+^ sAC^-/-^* (O) mice showing increased FM staining in *sAC^-/-^* mice. Arrows denote FM+ areas. P-O) Electron micrographs of mouse skin from *sAC^fl/fl^* (P) vs *sAC^-/-^* (Q) mice depicting keratinocytes showing an increase in highly melanized melanosomes in *sAC^-/-^* mice. Arrows denote stage IV melanosomes. All images are representative of N≥10 mice per cohort, mice combined over at least three litters.

To study the effects of sAC loss of function in melanocytes on epidermal pigmentation, we introduced the *K14::Kitl* allele which expresses stem cell factor (SCF, Kit ligand) in basal keratinocytes(*27*). Expression of SCF in basal keratinocytes recruits melanocytes to the basal cell layer thereby creating a “humanized” mouse model of epidermal pigmentation(*29*). We noted a clear increase in nail and epidermal pigmentation in *sAC^-/-^* compared to *sAC^fl/fl^* littermates (Fig. 4G-L), which was confirmed by measuring skin melanin content by HPLC (Fig. 4M). Histologic analysis of ear skin sections of *sAC^-/-^* versus *sAC^fl/fl^*littermates using Fontana-Masson showed enhanced melanin deposition in *sAC^-/-^*ears compared to *sAC^fl/fl^* ears (Fig. 4N-O). Consistent with our Fontana-Masson staining, ultrastructural examination revealed an increase in the number of highly-melanized melanosomes in the skin (Fig. 4P-Q). Further examination of ear and back skin revealed no differences in pigment enzyme expression between *sAC^-/-^*versus *sAC^fl/fl^* littermates (fig S7A) or epidermal melanocyte number (fig S7B-D). These data confirm that genetic sAC loss of function in *Tyr* expressing cells rescues ocular and epidermal pigmentation in OCA type 2 mice by increasing the activity of existing melanin synthetic machinery without altering melanocyte number or pigment gene expression.

### Pharmacologic sAC inhibition rescues ocular and cutaneous pigmentation in an OCA2 mouse model

Having observed multi-organ pigment increases due to pigment cell specific genetic sAC loss of function in an OCA type 2 mouse model, we reasoned that topical or systemic delivery of the previously described sACi (Fig. 2) would increase melanin synthesis in the eyes and skin of OCA type 2 mice. To assess ocular pigmentation, we delivered sACi intraperitoneally once a day for 30 days. Slit lamp examination and iris flat mount imaging revealed an increase in iris pigmentation in sACi treated versus vehicle treated mice (Fig. 5A-D). We recently developed a liquid chromatography mass spectroscopy (LC-MS)-based method that enables the measurement of melanin metabolites(*11, 16*). Applying this technique for the first time in live tissue, we assessed the effect of *ex vivo* melanin synthesis in the eye following sACi treatment. Compared to matched eyes from the same mouse, sAC inhibition *ex vivo* for 24 hours led to a significant increase in the levels of 5,6-dihydroxyindole-2-carboxylic acid (DHICA), a measure of eumelanin synthesis (Fig. 5E). Histochemical analysis demonstrated an increase in melanin in melanocytes in the iris and both pigmented epithelial cells and melanocytes in the ciliary body and choroid of the eye following sACi treatment (Fig. 5F-I). Thus, pharmacologic sAC inhibition rescues ocular pigmentation in OCA type 2 mice.

**Fig. 5.**
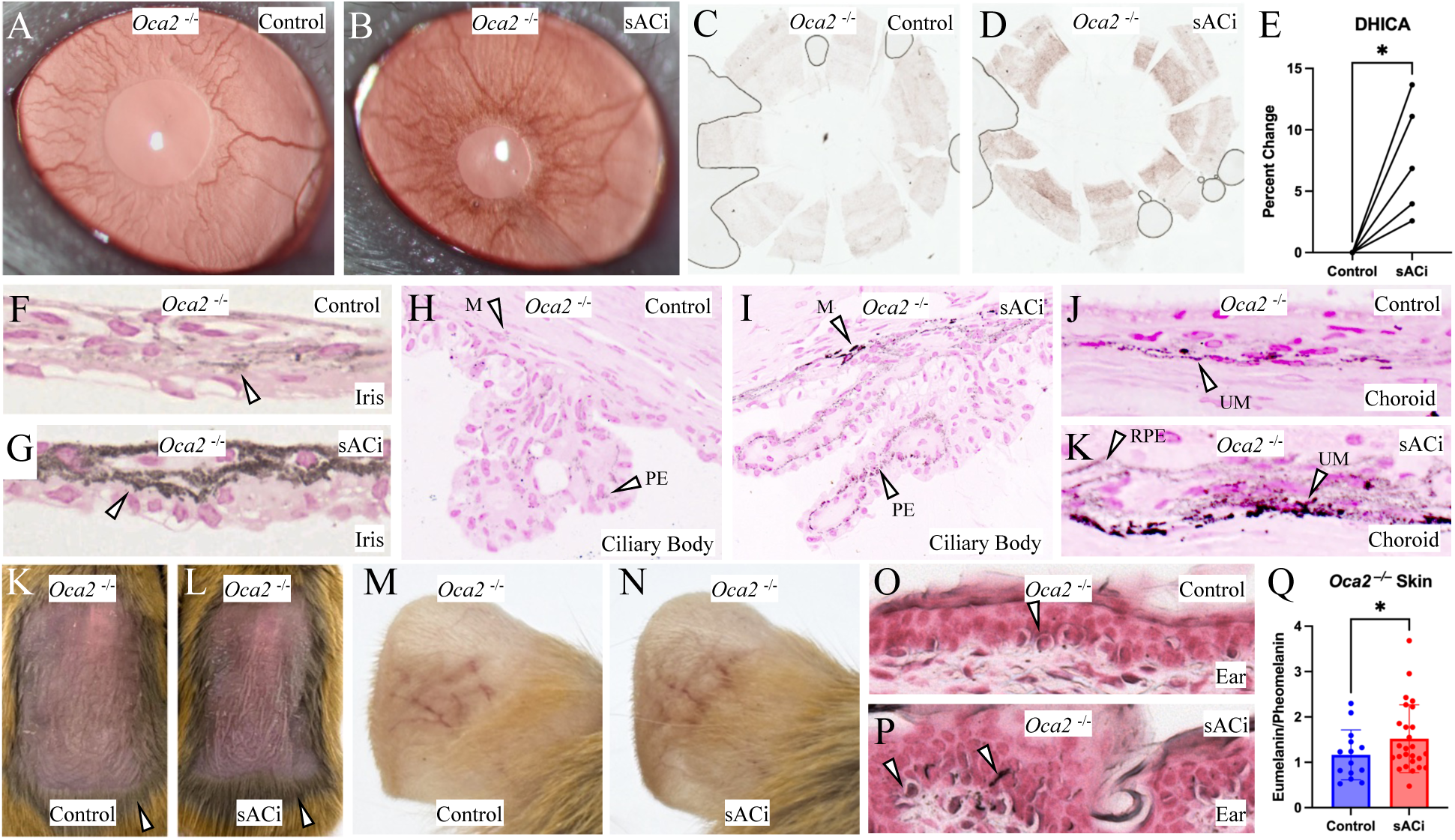
Pharmacologic sAC inhibition rescues ocular and cutaneous pigmentation in an OCA2 null mouse model. A-B) Iris slit lamp images following four weeks of intraperitoneal daily treatment of *Oca2^-/-^ sAC^fl/fl^* mice with control (DMSO, A) vs sACi (TDI-11155, B) showing darker iris pigmentation in sACi treated mice. C-D) Iris flat mount images from eyes treated as in (A, B) from control (DMSO, C) vs sACi (TDI-11155, D) mice showing darker iris tissue in sACi treated mice. E) LC-MS Q-TOF paired (left vs right) analysis of mouse eyes treated ex vivo with control (DMSO) or sACi (TDI-11155, 30µM) for 24 hours showing positive percent change in detected DHICA levels after treatment with sACi. N=5. F-G) Fontana-Masson (FM) staining of iris from mice treated as in (A, B) showing increased FM staining in the iris tissue in sACi treated mice (G) as compared to control (F). Arrows denote FM+ areas. H-I) FM staining of the ciliary body from mice treated as in (A, B) showing increased FM staining in both pigmented epithelial cells (see PE) and melanocytes (see M) in sACi treated mice (I) as compared to control (H). Arrows denote FM+ areas and source. J-K) FM staining of the choroid from mice treated as in (A, B) showing increased FM staining in RPE cells (see RPE) and uveal melanocytes (see UM) in sACi treated mice (K) as compared to vehicle (J). Arrows denote FM+ areas and source. K-N) Waxed skin (K-L) or ears (M-N) of *K14::Kitl^+^ Oca2^-/-^ sAC^fl/fl^* mice treated with control (DMSO, K, M) or sACi (TDI-11155, 1.5%, L, N) topically for 14 days showing darker skin and even darker adjacent hair (arrows) in sACi treated mice. O-P) FM staining of ears from mice treated as in (M-N) showing increased FM staining in ear skin in sACi treated (P) as compared to vehicle treated (O, control) mice. Arrows denote FM+ areas. Q) HPLC analysis of skin samples from *K14::Kitl^+^ Oca2^-/-^ sAC^fl/fl^* mice treated daily with control (DMSO) or sACi (TDI-11155, 1.5%) for 14 days showing elevated eumelanin/pheomelanin ratio in sACi treated skin samples. Welch’s t-test. *, P<0.05. Each point is one mouse. N≥14 per cohort. All images are representative of N≥10 mice per cohort, mice combined over at least three litters.

To examine the effects of sAC inhibition on epidermal pigmentation, we topically applied sACi daily for 14 days. Topical treatment of *K14::Kitl; Oca2^-/-^*mice with sACi increased epidermal pigmentation by visual inspection (Fig. 5K-N). Histologic analysis of melanin deposition using Fontana-Masson staining showed an increase in diffuse epidermal staining in skin treated with sACi compared to vehicle control (Fig. 5O-P). Moreover, HPLC analysis revealed that sAC inhibition significantly increased the eumelanin/pheomelanin ratio indicating improved production of eumelanin and decreased production of pheomelanin (Fig 5Q). As a control, we confirmed that sACi treatment of dorsal skin increases melanin synthesis without affecting pigment gene expression or melanocyte number (fig S8A-D). Thus, pharmacologic sAC inhibition is an effective method for rescuing ocular and epidermal pigmentation in OCA type 2 mice without altering melanocyte number or pigment gene expression.

## Discussion

OCA is a condition that predisposes affected individuals to a high lifetime risk of skin cancers, psycho-social stress, and visual impairment leading to significant disability. There are presently no FDA-approved therapies for OCA. Despite the significant burden associated with OCA, it has remained difficult to develop therapeutic options for this group of orphan conditions. Here, we demonstrate a novel therapeutic approach which aims to increase the activity of existing TYR by optimizing melanosome pH through modulation of the enzyme sAC.

Our approach is robustly supported by genetic evidence that modulation of sAC (*ADCY10*) activity influences human pigmentation. Our analysis of UK Biobank data showed that naturally-occurring deleterious variants in *ADCY10* shifted cutaneous pigmentation toward a darker phenotype and mitigated partial melanin-synthesis deficiencies. Notably, we conducted a phenome-wide analysis which revealed that rare *ADCY10* variants are largely unremarkable in their association with any specific phenotypic outcome(*30*), including male urological problems or hypercalciuria despite reported associations(*31*). As such, the limited systemic consequences of reduced sAC activity further reinforce its suitability as a therapeutic target for diseases of hypopigmentation.

Here, we show that sAC inhibition increases melanin synthesis under conditions of reduced TYR activity whether caused by acidic melanosome pH (OCA types 2 and 4) or by deleterious *TYR* variants (OCA type 1B). It is thought that melanosome pH alkalinizes during the process of melanosome genesis from about pH 5 in early-stage melanosomes to about pH 6.8 in late-stage melanosomes. As TYR activity is highest at near-neutral pH, it is probable that the increased pigmentation we observed in OCA type 1B models is due to alkalinization of early-stage melanosomes to a more neutral pH. This hypothesis is consistent with our ultrastructural observations that the distribution of OCA type 1B melanosomes shift toward more highly-melanized melanosomes due to sAC inhibition despite no observed changes in immunocytochemical markers of early- or late-stage melanosomes.

We also demonstrated that sAC inhibition alkalinizes melanosome pH and induces *de novo* melanin synthesis in mature human iRPE (figs. S1and S2E-F) (*26*). It is thought that the RPE contains lifelong melanin reserves which decline with age and in several degenerative conditions. Although the contribution of reduced RPE pigmentation to diseases, such as age-related macular degeneration (AMD), remains debated, lower pigment levels are associated with greater vulnerability to oxidative and light-induced stress. No approved therapy has been shown to restore melanin synthesis in the human RPE; thus, it has remained unclear whether mature RPE cells retain the capacity to generate new melanin. We now provide the first evidence that pharmacologic correction of RPE melanin deficits is achievable.

Our results demonstrate that alkalinizing melanosome pH is an effective method to rescue pigmentation in all pigmented cell types across all disease-relevant organ systems in both murine and human models of OCA. We observed that sACi treatment leads to a rapid increase in melanin synthesis without apparent signs of toxicity. Since loss of melanin synthesis in melanocytes and RPE cells likely accounts for most defects and disabilities observed in patients with OCA, these data suggest that sACi are a first-in-class approach for targeting melanosome pH to treat OCA. Moreover, we hypothesize that early treatment to increase pigmentation of the eye with sACi during critical periods of foveal development postnatally might be a viable therapeutic strategy for correcting some lifelong visual deficits in OCA.

Beyond individuals with OCA, hundreds of millions of people carry common genetic variants that similarly lower pigmentation by directly or indirectly reducing TYR activity. Our data suggest that sAC inhibition can enhance melanin synthesis across multiple genetic contexts, offering a potentially generalizable strategy for increasing pigmentation and decreasing skin cancer risk in a large portion of the population. Future work will focus on further development of lead drugs for ocular, cutaneous, and systemic delivery for the treatment of genetic and acquired disorders of pigmentation.

## Methods

### Genetic Association

Functional variants were defined as: (i) those labelled as a disease mutation (DM) in the Human Genetic Mutation Database (HGMD; 2024.1 version)(*32*); (ii) those predicted to be causing loss of function according to LOFTEE(*33*), and/or (iii) those with a CADD Phred-scaled score ≥20(*34*).

We then defined four key genetic groups: (A) individuals who carry no functional variants in the *OCA2* and *ADCY10* genes (baseline group); (B) individuals who carry ≥1 functional heterozygous variant in *ADCY10* only; (C) individuals who carry ≥1 functional heterozygous variant in *OCA2* only; and (D) individuals who carry ≥1 functional variant in each of *OCA2* and *ADCY10*.

We then binarized the UK Biobank(*35*) hair and skin color categories to include only the most ‘extreme’ values. For skin pigmentation, we classified participants with the darkest self-reported skin tone as ‘dark’ and those with the lightest self-reported tones as ‘light’. Similarly, for hair pigmentation, we examined individuals with the darkest (‘Black’) and lightest (‘Blonde’) self-reported hair colors, excluding intermediate shades.

After filtering the UK Biobank dataset to retain individuals meeting the predefined genotype combinations and selected pigmentation phenotypes, separate cohorts were constructed for the hair and skin pigmentation analyses. For the *OCA2* based analysis, the hair pigmentation cohort comprised 91,162 individuals (39,579 dark-haired and 51,583 light-haired). The corresponding skin pigmentation cohort included 40,952 individuals (3,724 dark-skinned and 37,228 light-skinned).

Associations between genotype group and pigmentation category were evaluated in R using binomial logistic regression, with pigmentation status specified as the binary outcome and genotype group as the predictor. In all models, “dark” pigmentation was defined as the reference outcome category and genotype group A as the reference group (such that odds ratios and 95% confidence intervals reflected the odds of light relative to dark pigmentation for groups B, C, and D compared with group A).

The same analytical framework was applied to the *TYR-*based analysis, substituting *OCA2* with *TYR* while retaining the same *ADCY10* grouping strategy, phenotype binarization, and statistical approach. In this analysis, the hair pigmentation cohort comprised 90,986 individuals (39,557 dark haired and 51,429 light haired), and the skin pigmentation cohort comprised 40,864 individuals (3,724 dark skinned and 37,140 light skinned).

### Mouse and human melanocyte culture

Mouse melanocytes (melan-p and melan-Ink4a/Arf^-/-^ [Ink4a]) were obtained from the Wellcome Trust Functional Genomics Cell Bank (St. George’s, University of London, UK) and cultured as previously published in RPMI 1640 supplemented with 10% FBS, 1% penicillin–streptomycin, 2 mM glutamine, and 200 nM 12-O-tetradecanoylphorbol-13-acetate (TPA) at 10% CO_2_ in a Thermo Scientific Heracell Vios 160i incubator (Yusupova et al., 2024, Hida et al., 2009, Sviderskaya et al. 2002). Human primary melanocytes containing biallelic variants in *OCA2,* NM_000275.3 [c.1327G<A, p.Val443Ile]; [c.1327G<A, p.Val443Ile] (p.V443I/p.V443I), NC_000015.10:g.[28017719_28020678delinsTTT]; [28017719_28020678delinsTTT] (D2.7kb/D2.7kb), and *TYR,* NM_000372.5 [c.980A>G, p.Tyr327Cys]; [c.929dup, p.Arg311Lysfs*7] (p.Y327C/p.R311X), and NM_000372.5 [.1118C>A, p.Thr373Lys]; [c.- 301C;575C>A;1205G>A] (p.T373K/p.[S192Y;R402Q]) were obtained from OCA type 2 and OCA type 1B presenting individuals, respectively, seen at the National Institutes of Health under NIH IRB approved protocol 09-HG-0035. Cells were cultured at 37 °C and 5% CO_2_ in a Thermo Scientific Heracell Vios 160i incubator in Ham’s F-10 (Gibco) supplemented with 5% fetal bovine serum (Corning), 2 mM L-Glutamine (Gibco), 250 ng/mL Amphotericin B (Gibco), 1% Penicillin Streptomycin (100 Units/mL and 100 μg/mL, respectively) (Gibco), 5 ng/mL basic fibroblast growth factor (Preprotech), 10 ng/mL Endothelin 1 (Sigma-Aldrich), 7.5 μg/mL 3-isobutyl-1-methylxanthine (Sigma-Aldrich), 30 ng/mL cholera toxin (Sigma-Aldrich), and 3.3 ng/mL Phorbol-12-myristate-13-acetate (Fisher Scientific). Media was replaced every third day. Mycoplasma testing was performed quarterly using commercial kits. Wild type and OCA2 null RPE cells were differentiated from iPSCs as previously described (*36*).

### *CRISPR KO of* Slc45a2

Ink4a melanocytes were transfected with mRNA expressing Cas9, mRNA expressing GFP, and three sgRNAs targeting *Slc45a2* (table S5). Forty-eight hours later single cells were sorted for GFP and cultured as single cell clones. Clones were tested for DNA deletion by PCR (table S5) and frame shift mutations by sequencing. Clones that passed this initial screen were grown and checked for loss of pigment production by visual inspection and *Slc45a2* mRNA by RT-qPCR (table S5).

### Phase microscopy and granule counting

Human primary OCA melanocytes cultured as described were treated with 0.1% DMSO with or without 30 μM TDI-11155 for 72 hours. Cells were assessed for the presence of darkly

pigmented granules after 72 hours using phase contrast microscopy with a 20X objective. Images were acquired using LAS X software (Leica) and processed for equal white balance in Adobe Lightroom (Adobe). Granules were counted using Aivia 14.1.0 “Cell Recipe” with “Enhance Bright Spots” background removal and vesicle partitioning.

### Measurement of melanin content by flow cytometry

Measurement of melanin by flow cytometry was performed according to the method previously published (*37*). OCA type 2 and OCA type 1B human melanocytes, melan*- Ink4a/Arf ^-/-^; Oca2 ^-/-^*and melan*-Ink4a/Arf ^-/-^; Slc45a2 ^-/-^* mouse melanocytes, and wild type and OCA2 null RPE cells were treated with normal culture media in appropriate CO_2_ and either control or 30 μM sAC inhibitor for the time described in the legend. Following treatment, cells were trypsinized and resuspended in 100 µL phosphate-buffered saline (PBS) without calcium and magnesium (Corning). Flow cytometry was performed using a BD Fortessa X20 5 laser analyzer. Data collection included forward scatter, side scatter, and scatter intensities at 355 nm. A minimum of 2000 events were collected and analyzed per replicate sample. All analysis was performed using FCS Express (DeNovo Software). Events were gated by side scatter and forward scatter to isolate live cells prior to further processing. Differential light scattering due to melanin was captured as mean scatter intensity through a 379/28 nm bandpass filter using a 355 nm ultraviolet laser for melanocytes and side-scatter at 400 nm for RPE cells. Mean scatter was statistically analyzed using unpaired Student’s t-test comparing vehicle to sAC inhibition treated cells.

### Cell and epidermal electron microscopy and quantitation

Human OCA type 2 and OCA type 1B melanocytes were grown to 50% confluence and treated for 72 hours with control or sAC inhibition. Following treatment, cells were washed in ice cold PBS and then fixed in 2.5% glutaraldehyde, 4% paraformaldehyde, 0.02% picric acid, 0.1 M sodium cacodylate buffer, washed three times with buffer, and post-fixed using 1% osmium tetroxide, 1.5% ferricyanide fixative overnight at room temperature for 60 minutes. Samples were then washed three times with buffer and stained with 1.5% uranyl acetate for 30 minutes at room temperature. Similar process was conducted on epidermal tissue from mice. Ocular samples were prepared as described (*36*). Tissues were rinsed in PBS, fixed in 2.5% glutaraldehyde, post-fixed, dehydrated through graded ethanol, and embedded in epoxy resin. Ultrathin sections on copper grids were stained with uranyl acetate and lead citrate and imaged on a JEOL JEM-1010 TEM. Following staining, samples were dehydrated by graded ethanol series and transitioned through acetonitrile. Cell monolayers were then embedded in Embed812 resin (Electron Microscopy Sciences). En face section measuring 55-60 nm cut using a Diatome diamond knife (Diatome) on a Leica Ultracut T ultramicrotome (Leica Microsystems) were contrasted with lead citrate and viewed on a JEM 1400 electron microscope (JEOL) operated at 100 kV. Digital images were captured on a Veleta 2K x 2K charge-coupled device camera (Olympus Soft Imaging Solutions). For cells in culture, melanosomes were staged and quantified manually in Adobe Illustrator (Adobe Inc., San Jose, CA, USA) using compiled images of 5 cells per treatment per cell line. Melanosomes were staged according to previously published standards for staging (*38*). In epidermal relative number of stage 4 melanosomes were noted.

### Mouse model including tamoxifen painting

The C3H/HeJ; *Tyr::Cre^ERT2^*; *Adcy10^fl/fl^*; *Oca2^p.R262X/p.R262X^* (*sAC*^fl/fl^; *Oca2*^-/-^) mouse line was generated according to methods previously described (*23, 27*). In brief, we introduced a previously characterized *Oca2* null allele (*28*) to homogeneity into our previously described *Tyr::Cre^ERT2^*; *sAC^fl/fl^* mice (*23, 27*) and then backcrossed this mouse onto a C3H/HeJ agouti mouse strain. *Tyr::Cre^ERT2^* and *sAC^fl/fl^*alleles were assessed using genomic PCR (fig. S4B and table S5). C3H/HeJ backcrossing was monitored using Transnetyx genetic monitoring service against a C3H/HeJ reference we provided. Transnetyx services were also used for monitoring *Oca2 R262X* genotype during backcrossing. The inducible knockout *sAC* allele was activated by painting the dorsum of all mice in the litter with 20 mM 4-hydroxytamoxifen (Sigma Aldrich H6278) on postnatal days 2, 3, and 4. Tamoxifen binds to the ERT2 protein fused to the Cre recombinase, thereby revealing a nuclear localization domain and allowing for Cre entry into the nucleus and gene recombination.

### Slit lamp examination

Gross morphology evaluation of the anterior of the eye was captured with a Nikon FS-3 Zoom Photo Slit Lamp equipped with a Canon EOS 60D digital camera. Briefly, mice were anesthetised with isoflurane using a rodent anaesthesia face mask (Braintree Scientific Inc.) and placed on a warm platform for slit lamp imaging. Eyes were kept moist with saline drops and additional lighting was provided by a dual gooseneck illuminator.

### Fundus imaging

The retina was evaluated with a Micron III retinal imaging microscope (Phoenix Research Labs). Mice were anesthetised with isoflurane using a rodent anaesthesia face mask (Braintree Scientific Inc.) and placed on a warm Micron Animal Stand which provides an articulatable platform for imaging. A small drop of 0.3% hypromellose lubricant eye gel (Genteal) was added at the time of imaging, acting as a coupling agent between the Micron III lens and the mouse eye, the gel also acts as a hydrating agent, preventing the mouse eye from drying.

### Histochemical and Fontana Masson staining of skin and eye tissue

After euthanasia, mouse ears and eyes were dissected and fixed in 10% neutral buffered formalin overnight at 4 °C. The tissues were rinsed in 1× PBS and transferred to 70% ethanol at 4 °C prior to paraffin embedding. Ears were placed in cassettes with sponges and processed using an automated tissue processor (70%, 85%, and 2×95% ethanol for 60 minutes each; 2×100% ethanol; 2× Histoclear for 60 minutes each; and 3× paraffin for 60 minutes each). The ears were embedded upright, and 7 µm-thick sections were obtained.

For the Fontana–Masson stain (Millipore Sigma HT200-1KT), sections were deparaffinized and rehydrated to deionized water. Freshly prepared ammoniacal silver solution was placed in a Coplin jar and heated to 58 °C. Slides were incubated in the solution for 30 minutes at 58 °C, until the tissue developed a yellowish-brown color, and then rinsed in several changes of deionized water. Sections were treated with 0.2% gold chloride solution for 30 seconds at room temperature, rinsed in deionized water, and incubated in 5% sodium thiosulfate solution for 2 minutes, followed by two additional rinses in deionized water. Sections were counterstained with Nuclear Fast Red for 5 minutes, rinsed, dehydrated through three changes of 100% ethanol, cleared with Histoclear, and mounted with mounting medium.

### Melanosome pH measurement

ARPE-19 cells were grown to a confluent monolayer. Cells were treated for four hours with the sAC inhibitor LRE1 (30μM) or vehicle (DMSO). Cells were then washed with their respective growth media, incubated with 10μM N-(3-((2,4-Dinitrophenyl) Amino)propyl)-N-(3-Aminopropyl) Methylamine, Dihydrochloride (DAMP) for 30 min, and fixed as previously described (*23*). Immunohistochemical labeling of DAMP and melanosomes was performed using goat polyclonal antibodies against dinitrophenol (anti-DNP, Oxford Biomedical Research, 1:200, catalog no. D04, lot no. d4.111212) and the mouse monoclonal antibody against human melanoma black 45 (HMB45) [Melanoma Marker Antibody (HMB45), Santa Cruz Biotechnology, 1:200, catalog no. sc-59305, lot nos. F1417, E1415], respectively. Immunohistochemistry and immunofluorescence quantitation of DAMP for relative pH measurements were then performed as described in (*23*) with the following change: images were acquired either using a Zeiss LSM 880. Fluorescent intensity for DNP staining was assessed only at all HMB45 + granules.

### Immunocytochemistry

Cells were cultured on sterile glass coverslips in 24-mm wells at 5.0 × 10^4^ cells per coverslip in their respective growth conditions for 24 hours. Cells were treated for 72 hours with 0.1% DMSO with or without 30µM TDI-11155, fixed for 15 minutes at room temperature in 2-4% paraformaldehyde, and rinsed with PBS. Immunocytochemical labelling of melanosomes was performed with HMB45 mouse mAb (Santa Cruz Biotechnology sc-59305) at 1:400 and TYRP1 mouse mAb (Abcam TA99 ab3312) at 1:100 in blocking buffer (PBS with 0.02% saponin, 1% BSA, and 2.5% normal goat serum) for one hour at room temperature. Samples were rinsed three times with blocking buffer and then incubated in donkey anti-mouse IgG (H+L) Alexa Fluor^TM^ 647 (ThermoFisher Scientific A-31571) at 1:300 in blocking buffer for 30 minutes. Cells were further rinsed twice more with blocking buffer and rinsed a final time in PBS before mounting in Prolong Gold Antifade with 4’,6-diamidino-2-phenylindole (DAPI) (Cell Signaling Technology, 8961S). Images were acquired using a Leica Stellaris 8 FALCON. HMB45 and TYRP1 positive granules were segregated by contrast threshold and quantified using Aivia 14.1.0 “Cell Recipe” with “Enhance Bright Spots” background removal and vesicle partitioning.

### qRT-PCR

RNA was generated from homogenized mouse eyes preserved in RNA*later* (ThermoFisher Scientific #AM7020) using the Qiagen RNAeasy RNA Purification Kit (Qiagen #74104). cDNA was then generated using the Applied Biosystems High-Capacity RNA-to-cDNA™ Kit (Thermo Fisher Scientific 4387406). Reverse transcription-quantitative PCR (qRT-PCR) was performed using the Applied Biosystems PowerUp™ SYBR™ Green Master Mix for qPCR (Thermo Fisher Scientific A25742) under standard protocol conditions using 10 μL reactions in an Applied Biosystems MicroAmp™ Optical 384-Well Reaction Plate with Barcode (Thermo Fisher Scientific 4309849). All reactions were run in a Quant Studio 6 (Thermo Fisher Scientific) and delta delta CT analysis was performed to determine relative transcription amongst samples normalized to *Gapdh* expression (table S5).

### Western Blot

Mouse back, ear, and eyes were homogenized in RIPA buffer with 1% protease/phosphatase inhibitor cocktail (Cell Signaling 5872S) on ice using a Dounce homogenizer. Lysates were prepared in Laemmli sample buffer with 2.5% β-mercaptoethanol. Proteins were separated using SDS-PAGE gel electrophoresis with a 1.5 mm, 8% polyacrylamide resolving gel. All gels were prepared and run using the Bio-RAD Mini-PROTEAN Tetra Gel system (Bio-RAD 1658000). Gels were run using a Bio-RAD Basic Power Supply (Bio-RAD 1645050) at 70 volts until the proteins had passed the stacking gel and 130 volts thereafter. Proteins were blotted onto Cytiva Amersham 0.45 NC Nitrocellulose membrane (Cytiva 10120-006) using the Bio-RAD Mini Trans-Blot system (Bio-RAD 1703935) run at 30 volts overnight at 4 degrees C. All blots were blocked in 5% milk-TBST and washed in TBST. Blots were probed with Vinculin XP Rabbit mAb (Cell Signaling 13901S) at 1:2000 in 5% BSA-TBST and Tyrosinase T311 Mouse mAb (Santa Cruz sc-20035) at 1:200 in 5% milk-TBST. HRP-linked Goat anti-Rabbit mAb (Cell Signaling 7074S) was used at 1:3000 for detection of vinculin while HRP-linked Goat-anti Mouse mAb H+L (Thermo Fisher Scientific G-21040) was used at 1:3000 for detection of tyrosinase. Pierce SuperSignal™ West Pico PLUS Chemiluminescent Substrate (Thermo Fisher Scientific 34580) was used to develop blots and blots were imaged using a Bio-Rad Gel Doc XR+ System (Bio-Rad, discontinued) with Bio-Rad Image Lab software (Bio-Rad version 6.1).

### LC-MS measurement of melanin metabolites

Liquid Chromatography-Mass Spectrometry (LC-MS) was performed according to the method previously published(*11*). Dulbecco’s Modification of Eagle’s Medium (DMEM) with 4.5 g/L glucose was prepared by The Media Preparation Core, Memorial Sloan Kettering Cancer Center, NY, US. DMEM medium was supplemented with 10% dialyzed FBS (Cytiva), 1% penicillin-streptomycin (Corning) and 0.11mM L-tyrosine. Mouse eyes were incubated in a 24-well tissue culture treated dish in DMEM media supplemented with either 0.1% DMSO or 30 μM TDI-11155. Eyes were quickly washed once with ice-cold PBS, then rinsed twice with ice-cold ddH_2_O. Metabolites were extracted using -80 °C 80:20 methanol: water (LC-MS grade methanol, Fisher Scientific). The tissue-methanol mixture was subjected to bead-beating for 90 seconds using a Tissuelyser cell disrupter (Qiagen). Extracts were centrifuged for 10 min at 13.2k rpm to pellet insoluble material and supernatants were transferred to clean tubes. The extraction procedure was repeated two additional times, and all three supernatants were pooled, dried in a speed-vac (Savant) and stored at -80 °C until analysis. The methanol-insoluble protein pellet was solubilized in 0.2 M NaOH at 95 °C for 20 min and quantified using the BioRad DC assay. On the day of metabolite analysis, dried cell extracts were reconstituted in 70% acetonitrile at a relative protein concentration of 2.5 mg/mL, and 8 µL of this reconstituted extract was injected for LC-MS-based targeted profiling. Cell extracts were analyzed by LC-MS as described previously (*39, 40*) using a platform comprised of an Agilent Model 1290 Infinity II liquid chromatography system coupled to an Agilent 6550 iFunnel time-of-flight mass spectrometer. Chromatography of metabolites utilized aqueous normal phase (ANP) chromatography on a Diamond Hydride column (Microsolv). Mobile phases consisted of: (A) 50% isopropanol, containing 0.025% acetic acid, and (B) 90% acetonitrile containing 5 mM ammonium acetate. To eliminate the interference of metal ions on chromatographic peak integrity and electrospray ionization, Ethylenediaminetetraacetic acid (EDTA) was added to the mobile phase at a final concentration of 6 µM. The following gradient was applied: 0-1.0 min, 0% B; 1.0-15.0 min, to 20% B; 15.0 to 29.0 min, 50% B; 29.1 to 37 min, 99% B. Raw data were analyzed using MassHunter Profinder 8.0 and MassProfiler Professional (MPP) 14.9.1 software (Agilent Technologies).

### HPLC melanin analysis

For melanin quantification, mouse tissue was homogenized and processed by alkaline hydrogen peroxide oxidation (AHPO) and hydroiodic acid (HI) hydrolysis according to the previously published method (*27, 41–43*). The degradation product pyrrole-2,3,5-tricarboxylic acid (PTCA) produced during AHPO was used to quantify eumelanin while the degradation product 4-amino-3-hydroxyphenylalanine (4-AHP) produced by HI hydrolysis was used to quantify pheomelanin. To convert the quantities of degradation products to melanin quantities, the amount of eumelanin (EM) was derived from the quantity of PTCA using the conversion factor 0.080, the quantity of benzothiazole-derived pheomelanin (BZ-PM) was derived from the quantity of TTCA using the conversion factor 0.034, and the amount of benzothiazine-derived pheomelanin (BT-PM) was derived from the quantity of 4-AHP using the conversion factor 0.007. Total pheomelanin (PM) was derived by summing the quantities of BZ-PM and BT-PM, and total melanin was derived by summing the quantities of PM and EM.

### *Tritium* in vivo *tyrosinase assay*

Tyrosinase activity of melanocytes in vivo was determined by measuring the amount of radioactive H_2_O produced from L-[Ring-3,5-^3^H]-Tyrosine, as previously described(*23*). Mouse melan-p2 melanocytes were incubated in six-well plates in media containing L-[Ring-3,5-^3^H]-Tyrosine (5 μCi/ml; PerkinElmer) for 4 or 8 hours in the presence or absence of LRE1 (30µM). Media (1.5 ml) from each well were removed and centrifuged at 1200 rpm in a microfuge (Eppendorf 5154 D) for 5 min. Supernatant (1 ml) was combined with 1 ml of 0.1 M citric acid containing 10% (w/v) activated charcoal to remove excess tyrosine and then centrifuged at 12,000 rpm for 5 min. ^3^H activity of the supernatant was determined using a scintillation counter. Media incubated in parallel wells containing no cells were used as a negative control for tyrosinase activity.

### Immunohistochemistry

Mice were waxed on their back seven days prior to tissue harvest to ensure synchronization of the hair cycling into the anagen hair stage. Back skin was harvested and fixed in 10% neutral buffered formalin overnight at 4 °C. On the following day, the fixative was replaced with 1X PBS to wash overnight followed by cryoprotection in 10% sucrose for at least 24 hours prior to embedding in the cryomedia and frozen. Cryosections were cut on a cryostat at 10µm thickness and placed at –80 °C until staining. The slides were left at room temperature to dry for two hours prior to staining. Slides were washed in TBS+0.1% Tween (TBST) with 0.3% Triton-X 100 and 0.1% glycine for 15 minutes rocking at room temperature. After 30 minutes of permeabilization, slides were washed twice in TBST for five minutes rocking at room temperature. Slides were stained with primary antibodies against DCT (TRP2-D-18, Santa Cruz Biotechnology sc-10451), SOX10 (Santa Cruz Biotechnology sc-17343), or TYRP1/PepI (*44*) diluted to 1:100-1:300 in 1X PIPES/HEPES/EGTA/Magnesium Sulphate Heptahydrate buffer (PHEM buffer (*45*)). Sections were incubated in primary antibody overnight at 4 °C in a humidified slide chamber. Slides were washed twice in TBST for 10 minutes rocking at room temperature prior to incubating for two hours in secondary antibody (donkey anti-goat IgG (H+L) Alexa Fluor^TM^ 488 [ThermoFisher Scientific A-11058], donkey anti-rabbit IgG (H+L) Alexa Fluor^TM^ 594 [ThermoFisher Scientific A-21207]) diluted to 1:1000 in TBST in humidified slide chambers at room temperature. Slides were washed twice in TBST for 20 minutes at room temperature, covered with Fluoromount-G mounting media, and coverslipped. Slides were allowed to set for 24 hours before analysis.

### Pigment Evaluation

Pigment in the anagen hair bulb was assessed in two ways using bright-field transmitted microscopy with a Nikon Eclipse 80i microscope. For the eumelanin to pheomelanin comparison, DAPI was used to evaluate normal hair bulb morphology prior to assessing hair color. Pheomelanin and eumelanin bulbs were distinguished based on their characteristic coloration. Pigmentation intensity was scored semi-quantitatively into five categories: no pigment, standard pheomelanin, high pheomelanin, low/mid eumelanin and high eumelanin. Between 30-80 hair bulbs were evaluated per sample across all animals and the observer was blinded to genotype. For pigment evaluation in DCT+ hair bulbs, the hair bulbs were evaluated for pigment in four categories: no eumelanin, low eumelanin, mid eumelanin and high eumelanin. For statistical analysis, bulbs were bucketed into no/low eumelanin and mid/high eumelanin. A total of 20 DCT+ hair bulbs were evaluated per animal and the observer was blinded to genotype.

### Image Acquisition and quantification for IHC

Images were acquired using a Nikon Eclipse 80i microscope at identical exposure and gain settings. For measuring % TYRP1 area in the interfollicular epidermis (IFE), TIFFs were processed in FIJI to split color channels to quantify the TYRP1 signal. The interfollicular epidermis (IFE) area was manually outlined using the freehand selection tool and added to ROI manager. Only the areas of IFE which were in focus were used for measuring TYRP1 % area. In the selected ROI, threshold was applied using the default method to account for true TYRPI signal and % area was measured. For quantifying SOX10 positive nuclei, hair bulbs were evaluated directly under the microscope and SOX10 positive nuclei counted manually. 14-23 hair bulbs were evaluated in each sample and normal hair bulb morphology was assessed using DAPI.

### Statistical analysis

All statistical analyses were performed using GraphPad Prism 10.0 (GraphPad Software). In most cases, comparison of means was performed using an unpaired, two-tailed t test (for two groups) with Welch’s correction or a one- or two-way ANOVA (for groups of three or more) with Tukey correction for multiple comparison and/or Welch’s correction for unequal variance. In cases where data was lognormally distributed, a two-tailed lognormal t-test was performed for two groups. Comparisons of median DAMP frequency distributions (nonparametric data) between conditions were analyzed using a Mann-Whitney *U* test.

## Acknowledgements

This research was conducted using the UK Biobank Resource under Application 53144, and we gratefully acknowledge the UK Biobank participants and coordinators. The authors thank all volunteers participating in UK Biobank and who have made this project possible. sAC inhibitors were a gift from Jochen Buck and Lonny R. Levin. We gratefully acknowledge Lee Cohen Gould, Juan Pablo Jimenez, Dena Almeida and the entire staff of the Electron Microscopy and Histology services of the Weill Cornell Medicine Microscopy and Image Analysis Core for their assistance in preparing and obtaining electron micrographs and preparation and staining of mouse skins for Fontana Masson pigment analysis.

## Authors contributions

S.J.G., R.S., P.I.S., M.H., B.P.B., and J.H.Z. designed the experiments. S.J.G., D.J.G., R.S., D.S.P., S.A., M.Y., J.B., Z.E., T.W., D.Z., J.Y., A.A., A.G., K.W., S.I., C.H., M.G.T., and P.I.S. conducted the experiments. S.J.G., S.A., D.Z., A.G., D.J.G., P.I.S., and J.H.Z. generated the figures. S.J.G., D.S.P., M.Y., A.G., J.Y., Q.C., S.S.G., S.K.L., and D.R.A. generated critical reagents. S.J.G., P.I.S., and J.H.Z. wrote the manuscript with all authors providing feedback.

## Data and materials availability

UK Biobank data are available under restricted access through a procedure described at http://www.ukbiobank.ac.uk/using-the-resource/. All other data supporting the UK Biobank findings of this study are available within the article. The scripts used to analyse the datasets included in this study are available at https://github.com/davidjohngreen/. All other data is either presented within the figures and tables or will be made available on public repositories.

## List of Supplementary Materials

Fig. S1. sACi treatment increases melanosome pH in RPE cells.

Fig. S2. sAC inhibition increases pigmented melanosomes and overall melanin synthesis in melanocytes and RPE cells with albinism mutations.

Fig. S3. sAC inhibitor treatment does not affect total melanosome numbers.

Fig. S4. Development of C3H/HeJ; *Tyr::Cre^ERT2^*; *Adcy10^fl/fl^*; *Oca2^p.R262X/p.R262X^* (*sAC*^fl/fl^; *Oca2*^-/-^) mice.

Fig. S5. Genetic sAC deletion in pigmented cells of the OCA type 2 eye increases pigment production without increasing overall melanosome genesis.

Fig. S6. Genetic loss of sAC induces increased eumelanin synthesis at the OCA type 2 hair bulb.

Fig S7. Genetic sAC deletion in pigmented cells of the skin does not affect melanogenesis related gene expression or melanocyte number.

Fig S8. sACi treatment increases melanin synthesis in the skin without affecting melanogenesis related gene expression or melanocyte number.

Table S1. Number of UK Biobank participants included in the hair and skin pigmentation analyses, stratified by *OCA2*–*ADCY10* genotype group

Table S2. Number of UK Biobank participants included in the hair and skin pigmentation analyses, stratified by *TYR*–*ADCY10* genotype group

Table S3. Association of *OCA2*–*ADCY10* genotype groups with hair and skin pigmentation traits.

Table S4. Association of *TYR*–*ADCY10* genotype groups with hair and skin pigmentation traits.

Table S5. Oligonucleotides used in the preparation and analysis of murine cell and animal models

## References

1. M. G. Thomas, J. Zippin, B. P. Brooks, in GeneReviews((R)), M. P. Adam et al., Eds. (Seattle (WA), 1993).

2. E. Z. Ma, A. E. Zhou, K. M. Hoegler, A. Khachemoune, Oculocutaneous albinism: epidemiology, genetics, skin manifestation, and psychosocial issues. Arch Dermatol Res 315, 107–116 (2023).

3. M. Kwa, M. Ravi, K. Elhage, L. Schultz, H. W. Lim, The risk of ultraviolet exposure for melanoma in Fitzpatrick skin types I-IV: A 20-year systematic review with meta-analysis for sunburns. J Eur Acad Dermatol Venereol 39, 1239–1253 (2025).

4. E. N. Grigoryan, Pigment Epithelia of the Eye: Cell-Type Conversion in Regeneration and Disease. Life (Basel) 12, (2022).

5. R. L. Mort, I. J. Jackson, E. E. Patton, The melanocyte lineage in development and disease. Development 142, 620–632 (2015).

6. P. Manga, S. Loftus, Genetics of Skin, Hair, and Eye Color in Human Pigmentation Disorders. Ann Hum Genet 89, 305–320 (2025).

7. N. W. Bellono, I. E. Escobar, E. Oancea, A melanosomal two-pore sodium channel regulates pigmentation. Scientific reports 6, 26570 (2016).

8. A. L. Ambrosio, J. A. Boyle, A. E. Aradi, K. A. Christian, S. M. Di Pietro, TPC2 controls pigmentation by regulating melanosome pH and size. Proc Natl Acad Sci U S A 113, 5622–5627 (2016).

9. L. Le et al., SLC45A2 protein stability and regulation of melanosome pH determine melanocyte pigmentation. Mol Biol Cell 31, 2687–2702 (2020).

10. J. Ancans et al., Melanosomal pH controls rate of melanogenesis, eumelanin/phaeomelanin ratio and melanosome maturation in melanocytes and melanoma cells. Exp Cell Res 268, 26–35 (2001).

11. Q. Chen et al., Measurement of Melanin Metabolism in Live Cells by [U-(13)C]-L-Tyrosine Fate Tracing Using Liquid Chromatography-Mass Spectrometry. J Invest Dermatol 141, 1810–1818 e1816 (2021).

12. F. Liu et al., Genetics of skin color variation in Europeans: genome-wide association studies with functional follow-up. Hum Genet 134, 823–835 (2015).

13. C. G. Summers et al., Does levodopa improve vision in albinism? Results of a randomized, controlled clinical trial. Clin Exp Ophthalmol 42, 713–721 (2014).

14. I. F. Onojafe et al., Nitisinone improves eye and skin pigmentation defects in a mouse model of oculocutaneous albinism. J Clin Invest 121, 3914–3923 (2011).

15. A. Gargiulo et al., AAV-mediated tyrosinase gene transfer restores melanogenesis and retinal function in a model of oculo-cutaneous albinism type I (OCA1). Mol Ther 17, 1347–1354 (2009).

16. D. Zhou et al., Two-pore channel 2 is required for soluble adenylyl cyclase-dependent regulation of melanosomal pH and melanin synthesis. Pigment Cell Melanoma Res, (2024).

17. Z. Li et al., Complete elucidation of the late steps of bafilomycin biosynthesis in Streptomyces lohii. J Biol Chem 292, 7095–7104 (2017).

18. M. Balbach et al., USPTO, Ed. (Cornell University and Tri-Institutional Therapeutics Discovery Institute, Inc, USA, 2022).

19. M. Balbach et al., On-demand male contraception via acute inhibition of soluble adenylyl cyclase. Nat Commun 14, 637 (2023).

20. M. Fushimi et al., Discovery of TDI-10229: A Potent and Orally Bioavailable Inhibitor of Soluble Adenylyl Cyclase (sAC, ADCY10). ACS Med Chem Lett 12, 1283–1287 (2021).

21. M. Miller et al., Design, Synthesis, and Pharmacological Evaluation of Second-Generation Soluble Adenylyl Cyclase (sAC, ADCY10) Inhibitors with Slow Dissociation Rates. J Med Chem 65, 15208–15226 (2022).

22. N. W. Bellono, I. E. Escobar, A. J. Lefkovith, M. S. Marks, E. Oancea, An intracellular anion channel critical for pigmentation. Elife 3, e04543 (2014).

23. D. Zhou et al., Mammalian pigmentation is regulated by a distinct cAMP-dependent mechanism that controls melanosome pH. Sci Signal. 11, eaau7987 (2018).

24. M. B. Dolinska et al., Ampyrone (4-Aminoantipyrine) is a Direct Agonist of Human Tyrosinase and Potential Therapeutic for Oculocutaneous Albinism and Disorders of Hypopigmentation. bioRxiv, 2025.2010.2013.682036 (2025).

25. A. George et al., TYROSINASE-Deficient Human Retinal Pigment Epithelium Exhibits Melanosome Maturation Defects. Invest Ophthalmol Vis Sci 66, 4 (2025).

26. K. J. Miyagishima et al., In Pursuit of Authenticity: Induced Pluripotent Stem Cell-Derived Retinal Pigment Epithelium for Clinical Applications. Stem Cells Transl Med 5, 1562–1574 (2016).

27. M. Yusupova et al., Distinct cAMP Signaling Microdomains Differentially Regulate Melanosomal pH and Pigmentation. J Invest Dermatol 143, 2019–2029 e2013 (2023).

28. H. Shoji et al., A nonsense nucleotide substitution in the oculocutaneous albinism II gene underlies the original pink-eyed dilution allele (Oca2(p)) in mice. Exp Anim 64, 171–179 (2015).

29. J. A. D’Orazio et al., Topical drug rescue strategy and skin protection based on the role of Mc1r in UV-induced tanning. Nature 443, 340–344 (2006).

30. A. C. f. G. Research. (Retrieved [03 03, 2026]).

31. A. Akbari et al., ADCY10 frameshift variant leading to severe recessive asthenozoospermia and segregating with absorptive hypercalciuria. Hum Reprod 34, 1155–1164 (2019).

32. P. D. Stenson et al., The Human Gene Mutation Database (HGMD((R))): optimizing its use in a clinical diagnostic or research setting. Hum Genet 139, 1197–1207 (2020).

33. K. J. Karczewski et al., The mutational constraint spectrum quantified from variation in 141,456 humans. Nature 581, 434–443 (2020).

34. M. Schubach, T. Maass, L. Nazaretyan, S. Roner, M. Kircher, CADD v1.7: using protein language models, regulatory CNNs and other nucleotide-level scores to improve genome-wide variant predictions. Nucleic Acids Res 52, D1143–D1154 (2024).

35. C. Bycroft et al., The UK Biobank resource with deep phenotyping and genomic data. Nature 562, 203–209 (2018).

36. A. George et al., In vitro disease modeling of oculocutaneous albinism type 1 and 2 using human induced pluripotent stem cell-derived retinal pigment epithelium. Stem Cell Reports 17, 173–186 (2022).

37. M. B. Dolinska et al., Ampyrone (4-Aminoantipyrine) is a Direct Agonist of Human Tyrosinase and Potential Therapeutic for Oculocutaneous Albinism and Disorders of Hypopigmentation. bioRxiv, (2025).

38. G. Raposo, M. S. Marks, Melanosomes--dark organelles enlighten endosomal membrane transport. Nat Rev Mol Cell Biol 8, 786–797 (2007).

39. Q. Chen et al., Rewiring of Glutamine Metabolism Is a Bioenergetic Adaptation of Human Cells with Mitochondrial DNA Mutations. Cell Metab 27, 1007–1025 e1005 (2018).

40. Q. Chen et al., Untargeted plasma metabolite profiling reveals the broad systemic consequences of xanthine oxidoreductase inactivation in mice. PLoS One 7, e37149 (2012).

41. S. Ito, S. Del Bino, T. Hirobe, K. Wakamatsu, Improved HPLC Conditions to Determine Eumelanin and Pheomelanin Contents in Biological Samples Using an Ion Pair Reagent. Int J Mol Sci 21, (2020).

42. S. Ito et al., Usefulness of alkaline hydrogen peroxide oxidation to analyze eumelanin and pheomelanin in various tissue samples: application to chemical analysis of human hair melanins. Pigment Cell Melanoma Res 24, 605–613 (2011).

43. K. Wakamatsu, S. Ito, Advanced chemical methods in melanin determination. Pigment Cell Res 15, 174–183 (2002).

44. V. Virador et al., Production of melanocyte-specific antibodies to human melanosomal proteins: expression patterns in normal human skin and in cutaneous pigmented lesions. Pigment Cell Res 14, 289–297 (2001).

45. M. Schliwa, K. Weber, K. R. Porter, Localization and organization of actin in melanophores. J Cell Biol 89, 267–275 (1981).

