## Supplementary Materials for "Targeting melanosome pH is an effective method for the treatment of oculocutaneous albinism"

**List of Supplementary Materials:**

Fig. S1. sACi treatment increases melanosome pH in RPE cells.

Fig. S2. sAC inhibition increases pigmented melanosomes and overall melanin synthesis in melanocytes and RPE cells with albinism mutations.

Fig. S3. sAC inhibitor treatment does not affect total melanosome numbers.

Fig. S4. Development of C3H/HeJ; *Tyr::Cre*$^{ERT2}$; *Adcy10*$^{fl/fl}$; *Oca2*$^{p.R262X/p.R262X}$ (*sAC*$^{fl/fl}$; *Oca2*$^{-/-}$) mice.

Fig. S5. Genetic sAC deletion in pigmented cells of the OCA type 2 eye increases pigment production without increasing overall melanosome genesis.

Fig. S6. Genetic loss of sAC induces increased eumelanin synthesis at the OCA type 2 hair bulb.

Fig S7. Genetic sAC deletion in pigmented cells of the skin does not affect melanogenesis related gene expression or melanocyte number.

Fig S8. sACi treatment increases melanin synthesis in the skin without affecting melanogenesis related gene expression or melanocyte number.

Table S1. Number of UK Biobank participants included in the hair and skin pigmentation analyses, stratified by *OCA2–ADCY10* genotype group

Table S2. Number of UK Biobank participants included in the hair and skin pigmentation analyses, stratified by *TYR–ADCY10* genotype group

Table S3. Association of *OCA2–ADCY10* genotype groups with hair and skin pigmentation traits.

Table S4. Association of *TYR–ADCY10* genotype groups with hair and skin pigmentation traits.

Table S5. Oligonucleotides used in the preparation and analysis of murine cell and animal models

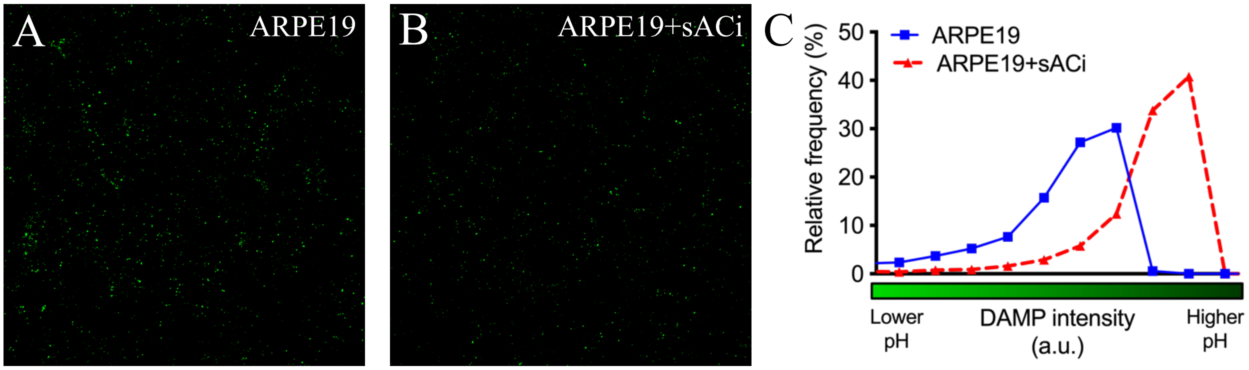


**Fig. S1. sACi treatment increases melanosome pH in RPE cells.** A) Human RPE cells (ARPE19) were grown to a confluent monolayer and treated with vehicle (A, DMSO, ARPE19) or sAC inhibitor (B, LRE1, 50µM, ARPE19+sACi) for fours hours and then treated with N-(3-((2,4-Dinitrophenyl) Amino)propyl)-N-(3-Aminopropyl) Methylamine, Dihydrochloride (DAMP). Cells were then fixed and immunostained for HMB45 and DAMP. Images show DAMP staining (green) only at HMB45 positive granules (melanosomes). C) Frequency distribution of relative intensity of DAMP across all melanosomes (>1000) in cells (>20 per coverslip) imaged from two separate coverslip showing that sACi alkalinizes melanosome pH in human RPE cells. Representative experiment performed three times.

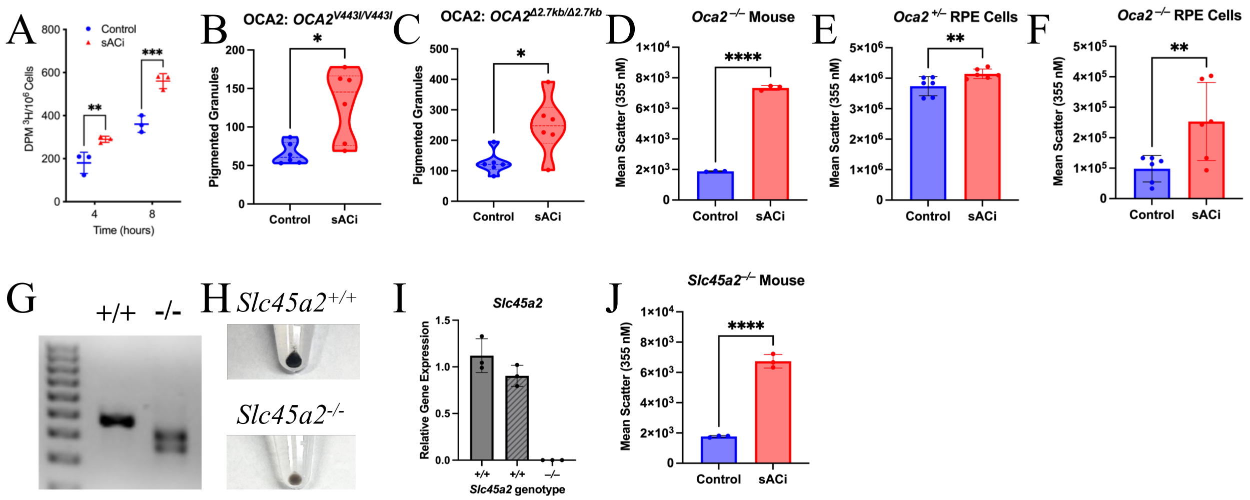


**Fig. S2. sAC inhibition increases pigmented melanosomes and overall melanin synthesis in melanocytes and RPE cells with albinism mutations.** A) Melan-p2 cells treated with ${}^3$H-H$_2$0 and either vehicle (DMSO) or sACi (LRE1, 50μM) for the time indicated. Media was collected and subjected to column chromatography (technical duplicates) as previously reported and ${}^3$H-H$_2$0 levels were measured. Cells were counted and used for normalization. Three distinct wells of cells were analyzed per experiment. Representative experiment performed three times. B-C) Human melanocytes derived from patients with OCA2 treated with control (DMSO) or sACi (TDI-11155, 30μM) for 24 hours and photographed by phase microscopy. Melanized melanosomes were counted per high powered field (3-5 cells). Each field is represented by a single point on the graph. Multiple high-powered fields were imaged across multiple (N>3) replicates. D-F) Melanin content measured by side scattered on flow cytometry in OCA2 KO Ink4a mouse melanocytes (D) and human OCA2 $^{+/-}$ (het) (E) and human OCA2$^{-/-}$ (F) RPE cells treated for 24 hours (D) or two weeks (E-F) with either control (DMSO) or sACi (TDI-11155, 30μM). G) PCR of Slc45a2 gene showing CRIPSR KO of genetic material confirmed to generate frame shift mutation. N=6 per condition. H) Cell pellets of *Slc45a2*$^{+/+}$ and CRISPR KO *Slc45a2*$^{-/-}$ Ink4a melanocytes. I) qRT-PCR of Slc45a2 in two wild-type Ink4a clones (+/+) and the Slc45a2 KO (-/-) clone. J) Melanin content measured by side scattered on flow cytometry in SLC45A2 KO Ink4a mouse melanocytes treated for 24 hours with either control (DMSO) or sACi (TDI-11155, 30μM). N=3. Two-way ANOVA with multiple comparison (A), Welch's t-test (B-C, G), or student's t-test (D-F). *,P<0.05; **,P<0.01; ***,P<0.001; and ****,P<0.0001. Each point is one mouse or biological replicate.

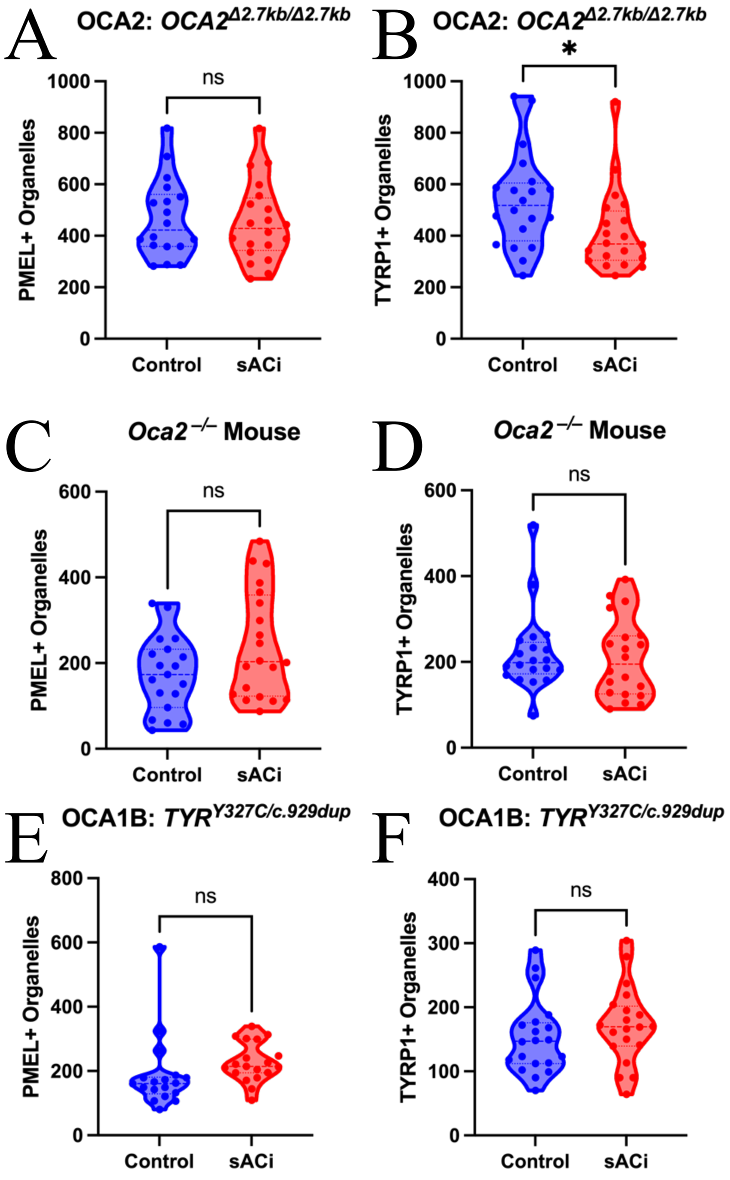


**Fig. S3. sAC inhibitor treatment does not affect total melanosome numbers.** Human and mouse melanocytes were treated with control (DMSO) or sACi (TDI-11155, 30µM) for 72 hours and then immunostained for PMEL or TYRP1. A-B) Human OCA2 null melanocytes (delta2.7kb/delta2.7kb) immunostained for PMEL (HMB45, A) or TYRP1 (TA-99, B). C-D) Mouse OCA2 null melanocytes generated by CRISPR KO immunostained for PMEL (HMB45, C) or TYRP1 (TA-99, D). E-F) Human OCA1B melanocytes (Y327C/c.929dup / Y327C/c.929dup) immunostained for PMEL (HMB45, E) or TYRP1 (TA-99, F). For all immunostaining total organelle number was quantified per cell. Welch's t-test. *,P<0.05. ns=not significant. Each point is one cell. Graphs are combined data from three experiments performed with duplicate coverslips per condition.

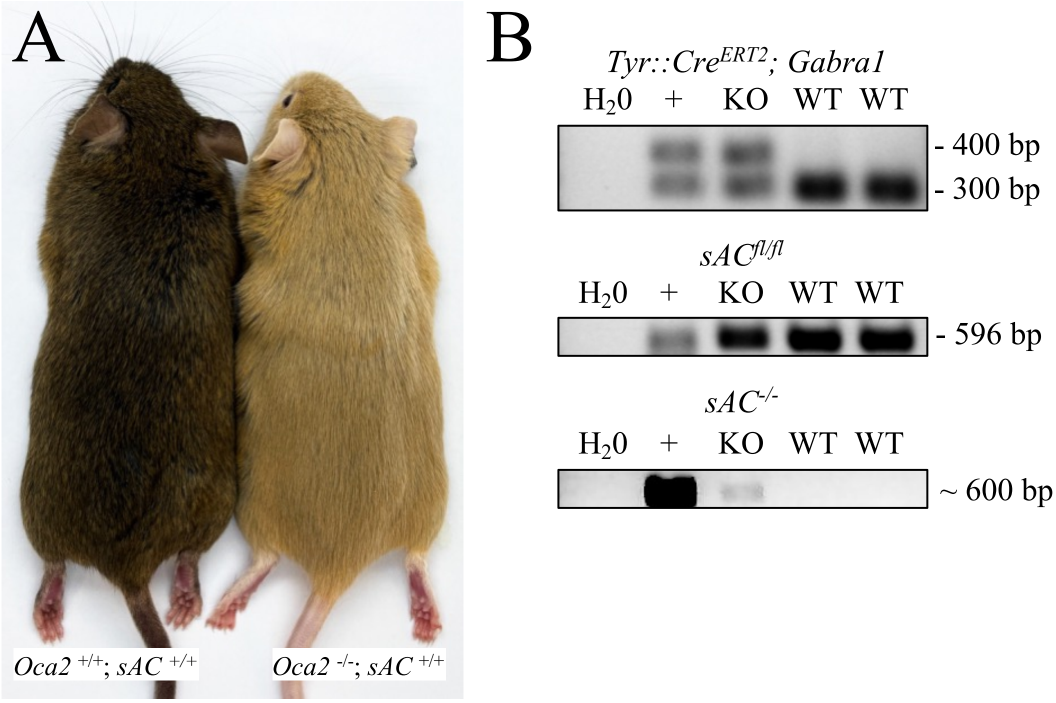


**Fig. S4. Development of C3H/HeJ; *Tyr::Cre*$^{ERT2}$; *Adcy10*$^{fl/fl}$; *Oca2*$^{p.R262X/p.R262X}$ (*sAC*$^{fl/fl}$*; OCA2*$^{-/-}$) mice.** A) Image of C3H/HeJ *Oca*$^{+/+}$*;sAC*$^{+/+}$ mice and C3H/HeJ *Oca*$^{-/-}$*;sAC*$^{+/+}$ mice showing depigmentation of wildtype *C3H/HeJ* following introduction of *Oca*$^{-/-}$ allele. B) Genomic PCR demonstrating genotyping method for C3H/HeJ *Oca*$^{-/-}$*;sAC*$^{fl/fl}$ (WT) and C3H/HeJ *Oca*$^{-/-}$*;sAC*$^{-/-}$ (KO) mice showing PCR results for *Tyr::Cre*$^{ERT2}$ (400 bp)*; Gabra1* (300 bp, PCR control), *sAC*$^{fl/fl}$*,* and *sAC*$^{-/-}$. Samples shown are water control ($H_20$), positive control (+), sAC$^{-/-}$ (KO) and sAC$^{fl/fl}$ (WT) mice. *sAC*$^{fl/fl}$ allele maintained in KO due to non-melanocyte DNA. Presence of ~600 bp band in *sAC*$^{-/-}$ samples confirms KO.

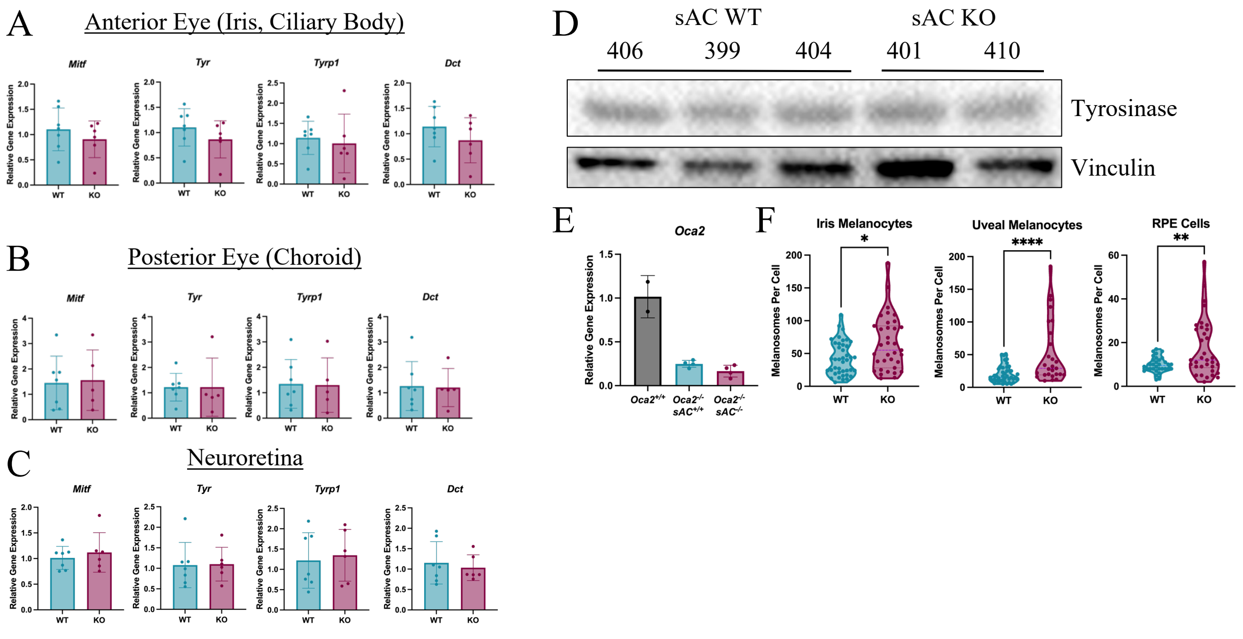


**Fig. S5. Genetic sAC deletion in pigmented cells of the OCA type 2 eye increases pigment production without increasing overall melanosome genesis.** A-C) qRT-PCR of *Mitf*, *Tyr*, *Tyrp1*, and *Dct/Tyrp2* from microdissections of anterior (A), posterior (B), and neuroretina (C) portions of the eye from $sAC^{fl/fl}$ (WT) vs $sAC^{-/-}$ (KO) OCA2 mice. Each point is one mouse. Each point is one mouse. N≥6. D) Western blot of whole eye tissue for TYR (T311) in $sAC^{fl/fl}$ (sAC WT) vs $sAC^{-/-}$ (sAC KO) OCA2 mice. 399, 401, 404, 406, and 410 are mouse numbers. E) qRT-PCR of *Oca2* expression in the eye in wildtype mice ($Oca2^{+/+}$) compared to $Oca2^{-/-}$ mice with either wildtype or KO sAC genotype. Each point is one mouse. F) Combined stage 3 and 4 melanosomes on electron micrographs of iris and uveal melanocytes and RPE cells in the eyes of sAC WT and sAC KO OCA2 mice. Each point represents one cell. Combined quantitation over N≥5 mice per genotype. Lognormal t-test (iris and uveal melanocytes) or Welch's t-test (RPE cells), *,P<0.05; **,P<0.01; ***,P<0.001; and ****,P<0.0001. Each point is one cell.

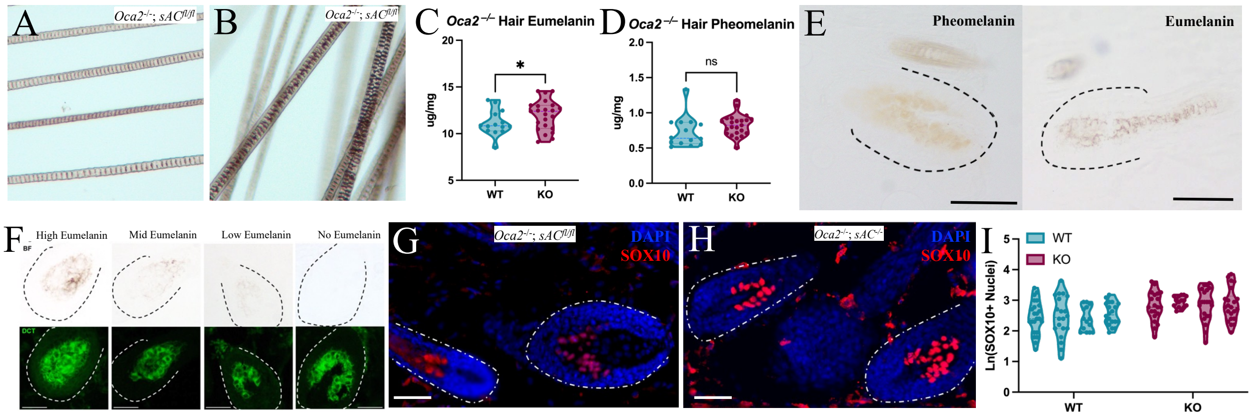


**Fig. S6. Genetic loss of sAC induces increased eumelanin synthesis at the OCA type 2 hair bulb.** A-B) Representative hair banding from $sAC^{fl/fl}$ vs $sAC^{-/-}$ mice. C) Eumelanin levels measured by HPLC in hair from $sAC^{fl/fl}$ (WT) vs $sAC^{-/-}$ (KO) OCA2 mice. N ≥12 per cohort. D) Pheomelanin levels measured by HPLC in hair from $sAC^{fl/fl}$ (WT) vs $sAC^{-/-}$ (KO) OCA2 mice. N≥12 per cohort. E) Example images of pheomelanin (left) and eumelanin (right) hair follicles. F) Representative examples of DCT+ (green) hair bulbs showing a range of measured eumelanin levels by bright field in OCA2 mice. G-H) Sox10 (red) immunostained and DAPI (blue) stained representative section showing total melanocyte number in hair bulbs from $sAC^{fl/fl}$ vs $sAC^{-/-}$ mice. I) Quantitation of Sox10 positive nuclei per hair bulb in $sAC^{fl/fl}$ (WT) vs $sAC^{-/-}$ (KO) mice. Welch's t-test. *, $P<0.05$. ns=not significant. C-D) Each point is one mouse. I) Each violin plot represents one mouse and each point represents one section. Scale bar = 100µm.

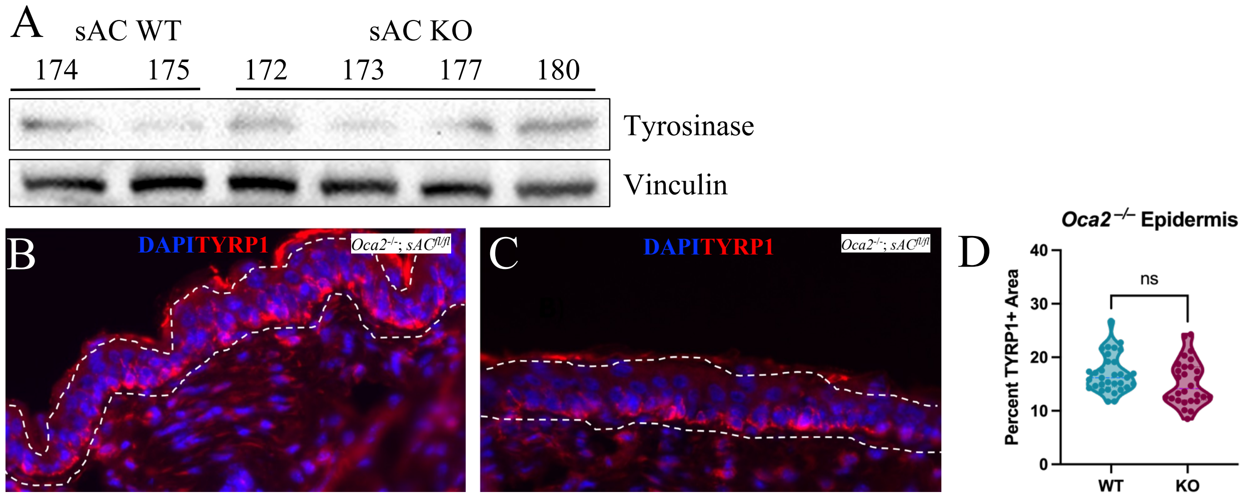


**Fig S7. Genetic sAC deletion in pigmented cells of the skin does not affect melanogenesis related gene expression or melanocyte number.** A) Western blot analyses of tyrosinase (T311) levels in skin of *K14::SCF*$^{+}$ *sAC*$^{fl/fl}$ (sAC WT) vs *sAC*$^{-/-}$ (sAC KO) OCA2 mice. Vinculin serves as loading control. B-C) TYRP1 immunostaining and DAPI staining of dorsal skin from *K14::SCF*$^{+}$ *sAC*$^{fl/fl}$ (B) and *sAC*$^{-/-}$ (C) mice. D) Percent of TYRP1+ area of the OCA2 epidermis as a proxy for melanocyte cell number from mice as described in (B-C). Welch's t-test. ns=not significant. Each point represents one image. At least 8 images were quantified per animal, and 3 animals were quantified per genotype.

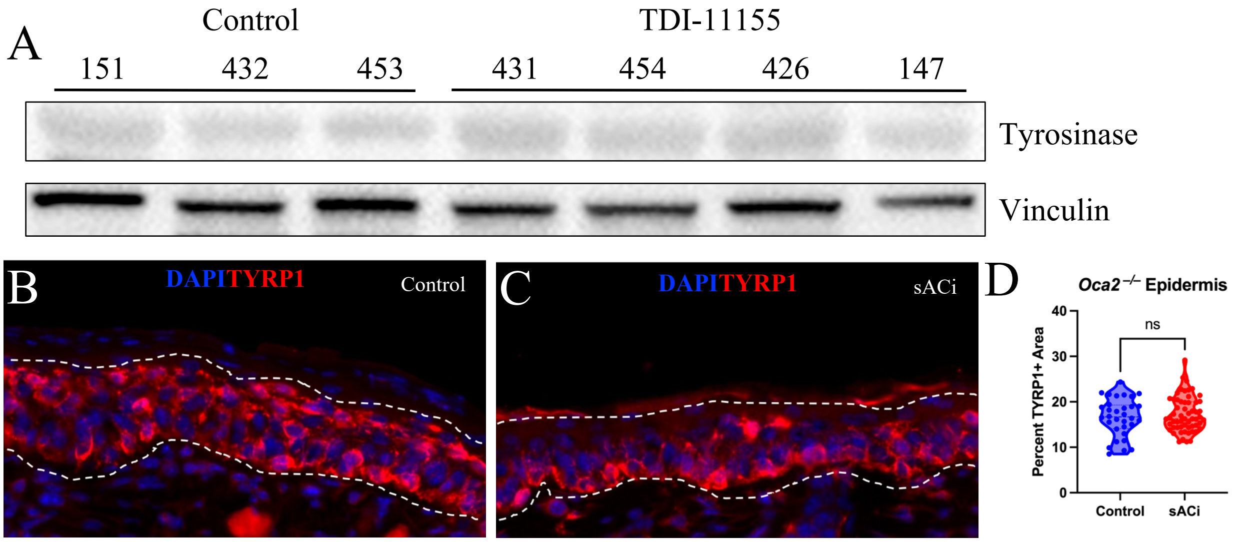


**Fig S8. sACi treatment increases melanin synthesis in the skin without affecting melanogenesis related gene expression or melanocyte number.** A) Tyrosinase (T311) Western blot analysis from *K14::SCF*$^{+}$ *sAC*$^{fl/fl}$ OCA2 mouse skin treated with either DMSO (control) or 30 mM TDI-11155 sACi for 14 days. B-C) TYRP1 immunostaining and DAPI staining of dorsal skin from *K14::SCF*$^{+}$ *sAC*$^{fl/fl}$ OCA2 mice treated with Control (B, DMSO) or sACi (C, 30 mM TDI-11155) for 14 days showing TYRP1+ epidermis. D) Quantitation of TYRP+ area as a proxy for melanocyte cell number. Welch's t-test. ns=not significant. Each point represents one image. At least 3 images were quantified per animal, and at least 7 animals were quantified per condition.

**Table S1.** Number of UK Biobank participants included in the hair and skin pigmentation analyses, stratified by *OCA2–ADCY10* genotype group

| Genotype group | Hair | | Skin | |
|---|---|---|---|---|
| | Dark | Light | Dark | Light |
| *OCA2* (ref) · *ADCY10* (ref) | 37,438 | 48,461 | 3,372 | 34,989 |
| *OCA2* (ref) · *ADCY10* (↓) | 529 | 119 | 208 | 83 |
| *OCA2* (↓) · *ADCY10* (ref) | 1,586 | 2,992 | 134 | 2,151 |
| *OCA2* (↓) · *ADCY10* (↓) | 26 | 11 | 10 | 5 |

↓ denotes reduced gene function due to the presence of ≥1 predicted deleterious variant; "ref" denotes absence of a predicted deleterious variant. Predicted deleterious variants were defined as: (i) those labelled as a disease mutation (DM) in the Human Genetic Mutation Database (HGMD; 2024.1 version (ii) those predicted to cause loss of function according to LOFTEE, and/or (iii) those with a CADD Phred-scaled score ≥20

**Table S2.** Number of UK Biobank participants included in the hair and skin pigmentation analyses, stratified by *TYR–ADCY10* genotype group

| Genotype group | Hair | | Skin | |
|---|---|---|---|---|
| | Dark | Light | Dark | Light |
| *TYR* (ref) · *ADCY10* (ref) | 37,856 | 48,025 | 3,464 | 34,655 |
| *TYR* (ref) · *ADCY10* (↓) | 542 | 126 | 213 | 80 |
| *TYR* (↓) · *ADCY10* (ref) | 1,145 | 3,274 | 42 | 2,397 |
| *TYR* (↓) · *ADCY10* (↓) | 14 | 4 | 5 | 8 |

↓ denotes reduced gene function due to the presence of ≥1 predicted deleterious variant; "ref" denotes absence of a predicted deleterious variant. Predicted deleterious variants were defined as: (i) those labelled as a disease mutation (DM) in the Human Genetic Mutation Database (HGMD; 2024.1 version (ii) those predicted to cause loss of function according to LOFTEE, and/or (iii) those with a CADD Phred-scaled score ≥20

**Table S3.** Association of *OCA2–ADCY10* genotype groups with hair and skin pigmentation traits.

| Genotype group | Hair | | Skin | |
|---|---|---|---|---|
| | Odds ratio (95% CI) | P-value | Odds ratio (95% CI) | P-value |
| *OCA2* (ref) · *ADCY10* (ref) | 1.00 (reference) | not applicable | 1.00 (reference) | not applicable |
| *OCA2* (ref) · *ADCY10* (↓) | 0.17 (0.14, 0.21) | $< 2.2 \times 10^{-16}$ | 0.04 (0.03, 0.05) | $< 2.2 \times 10^{-16}$ |
| *OCA2* (↓) · *ADCY10* (ref) | 1.46 (1.37, 1.55) | $< 2.2 \times 10^{-16}$ | 1.55 (1.30, 1.86) | $1.56 \times 10^{-6}$ |
| *OCA2* (↓) · *ADCY10* (↓) | 0.33 (0.15, 0.64) | $1.88 \times 10^{-3}$ | 0.05 (0.02, 0.14) | $3.13 \times 10^{-8}$ |

Odds ratios (ORs) are shown with 95% confidence intervals (CIs) from binomial logistic regression models. Analyses were performed separately for hair and skin pigmentation traits and stratified by *OCA2–ADCY10* genotype group.

↓ denotes reduced gene function due to the presence of ≥1 predicted deleterious variant; “ref” denotes absence of a predicted deleterious variant. For more information see the Methods section and Figure 1.

**Table S4.** Association of *TYR–ADCY10* genotype groups with hair and skin pigmentation traits.

| Genotype group | Hair | | Skin | |
|---|---|---|---|---|
| | Odds ratio (95% CI) | P-value | Odds ratio (95% CI) | P-value |
| *TYR* (ref) · *ADCY10* (ref) | 1.00 (reference) | not applicable | 1.00 (reference) | not applicable |
| *TYR* (ref) · *ADCY10* (↓) | 0.18 (0.15, 0.22) | $< 2.2 \times 10^{-16}$ | 0.04 (0.03, 0.05) | $< 2.2 \times 10^{-16}$ |
| *TYR* (↓) · *ADCY10* (ref) | 2.25 (2.10, 2.41) | $< 2.2 \times 10^{-16}$ | 5.68 (4.24, 7.85) | $< 2.2 \times 10^{-16}$ |
| *TYR* (↓) · *ADCY10* (↓) | 0.23 (0.06, 0.65) | 0.00466 | 0.16 (0.05, 0.54) | $4.84 \times 10^{-03}$ |

Odds ratios (ORs) are shown with 95% confidence intervals (CIs) from binomial logistic regression models. Analyses were performed separately for hair and skin pigmentation traits and stratified by *TYR–ADCY10* genotype group.

↓ denotes reduced gene function due to the presence of ≥1 predicted deleterious variant; "ref" denotes absence of a predicted deleterious variant. For more information see the Methods section and Figure 1.

**Table S5.** Oligonucleotides used in the preparation and analysis of murine cell and animal models

| Gene Target | Oligonucleotide Sequences |
|---|---|
| **Genotyping Primers for Mouse Models** | |
| *Adcy10*$^{-/-}$ (*sAC*$^{-/-}$) | F: 5'-GGACAGAAAGTAGAATGACTATCCCCCATTG-3'<br>R: 5'-CCGCTCACCTCTTTTCGGATTACATC-3' |
| *Adcy10*$^{fl/fl}$ (*sAC*$^{fl/fl}$) | F: 5'-GGACAGAAAGTAGAATGACTATCCCCCATTG-3'<br>R: 5'-AACCAAATCCCTAATGCCGACATGC-3' |
| *Tyr::Cre*$^{ERT2}$ | F: 5'-CCTGGAAAATGCTTCTGTCCG-3'<br>R: 5'-CAGGGTGTTATAAGCAATCCC-3' |
| *Gabra1* (PCR control) | F: 5'-CAATGGTAGGCTCACTCTGGGAGATGATA-3'<br>R: 5'-AACACACACTGGCAGGACTGGCTAGG-3' |
| **CRISPR KO sgRNAs, Genomic PCR, and Sequencing Primers** | |
| *Slc45a2* (CRISPR KO) | sgRNA 1: 5'-UGAUAGGUAUGGGUGUCAGU-3'<br>sgRNA 2: 5'-AGUCUCCCUGUGGAUCUCUU-3'<br>sgRNA 3: 5'-CCACCGCAUAGCAAAACUCU-3'<br>F: 5'-CCCCTTTGCTTCTCACCGTA-3'<br>R: 5'-CAGGATGTATGGTCTCCGGC-3'<br>Sanger Sequencing: 5'-GATGTATGGTCTCCGGCGAC-3' |
| **qRT-PCR Primers** | |
| *Gapdh* | F: 5'- GTGGAGTCATACTGGAACATGTAG-3'<br>R: 5'- AATGGTGAAGGTCGGTGTG-3' |
| *Mitf* | F: 5'-CTGGGTTTCTCCTGATTGGTC-3'<br>R: 5'-TTGTCCTTTTTCTGCCTCTCT-3' |

| | |
|---|---|
| *Tyr* | F: 5’-TAACTTACTCAGCCCAGCATC-3’<br>R: 5’-CGTAATAGTGGTCCCTCAGGT-3’ |
| *Tyrp1* | F: 5’-AATGGAGAGTGGTCTGTGAATC-3’<br>R: 5’-CTGGGTTTCTCCTGATTGGTC-3’ |
| *Dct* | F: 5’-ATGACCCAACGCTGATTAGTC-3’<br>R: 5’-CCTTCATAGGTTCCATTACACAGT-3’ |
| *Slc45a2* | F: 5’-GATTGCCCCTTTGGCTCTGT-3’<br>R: 5’-CCAAGATGGGGCTTAGGAGC-3’ |
